# A neural geometry for forelimb proprioception in the cervical spinal cord

**DOI:** 10.1101/2025.09.26.678887

**Authors:** Tejapratap Bollu, Martyn Goulding

## Abstract

Precise, real-time somatosensory feedback is essential for coordinated movement, yet how the nervous system constructs a kinematic representation of the limb from this feedback is unclear. Here we show that neurons in the cervical spinal cord generate a precise, ordered, low-dimensional representation of the forelimb’s kinematic state from muscle and tendon afferent input. This representation arises from two classes of molecularly distinct, developmentally defined dorsal horn neurons. One encodes limb speed, position and direction. The other exhibits phasic, precise, or radially-periodic firing fields that tile kinematic space. Transient perturbations of muscle and tendon afferents in mice reaching to spatial targets cause end-point errors as predicted by deficits in the neural code. These findings show that spinal networks, one synapse removed from the periphery, are sufficient to construct a precise representation of forelimb movement.

## Main Text

How the nervous system generates a representation of the body in time and space is for the most part poorly understood. Body representations are thought to emerge from proprioception(*1*, *2*) - the sense of self that animals use to perceive the position, movement, and force of their body parts - by the integrating of sensory input from muscles, joints, and skin to construct an internal map of the body’s posture and configuration in space(*3*). Although fully-spinalized vertebrates can account for their body schema during goal directed movements(*4–6*), neurophysiological descriptions of these body representations have been largely limited to the brain(*7–12*). By contrast, little is known about the how this information is encoded at spinal levels. The anatomical organization of sensory inputs to the spinal cord suggests that it has the capacity to integrate information across various muscle groups and skin territories to build a kinematic representation of the body(*13–15*). Using *in vivo* electrophysiological recordings from the cervical spinal cords of awake and anesthetized mice, we have discovered that the spinal cord relies on molecularly distinct, developmentally defined neural classes to construct an ordered low-dimensional representation of forelimb kinematics using information derived from muscle and tendon sensory afferents.

### The cervical spinal cord contains a precise representation of forelimb kinematics

Recording single neurons in the cervical spinal cord during forelimb movement has proved to be intractable due to the motion of the cord relative to the probe, thus limiting experiments to wrist movements where the limb is restrained (*16*). To overcome this issue, we developed a novel spinal device that stabilizes the cord (Fig. S1A) and recorded from the medial deep dorsal horn of the cervical spinal cord of awake mice while using key-point tracking (*17*) to capture the kinematics of spontaneous forelimb movements (Fig 1A). This area is densely innervated by muscle and tendon sensory afferents(*18*), making it a likely target for neurons encoding spatial sensory information. Recordings contained well isolated single units with high signal-to-noise ratio (Fig S1B), enabling us to identify the neural correlates of hand kinematics. Encoding models, ranging from linear to flexible nonlinear, were used to estimate how much kinematic information these neurons carry (Fig. 1B; Methods). The flexible models (Recurrent Neural Networks & Gradient Boosting Trees, Methods) we used made few assumptions and thus set an upper bound on the recoverable information. A non-linear interpretable additive model (Generalized Additive Model, Methods) enabled us to directly estimate the contributions of individual kinematic parameters. Most neurons (37/42, from 2 mice) carried significant kinematic information that was dominated by position rather than velocity or acceleration (Fig 1C). Their activity was diverse, with some neurons firing tonically and others sparsely and in bursts (Fig S1C). Tuning for kinematic parameters varied across neurons (Fig S1D-I), with neurons showing U-shaped tuning (Fig. S1E), sparse spatially restricted activity (Fig. S1G) or monotonic increases (Fig. S1I). These results established that structured neural representations for hand kinematics exist in the spinal cord, but they did not reveal the underlying sensory code which requires separating the sensory component of neural representations from confounds including intrinsic dynamics, descending modulation, and premotor commands.

**Fig. 1.**
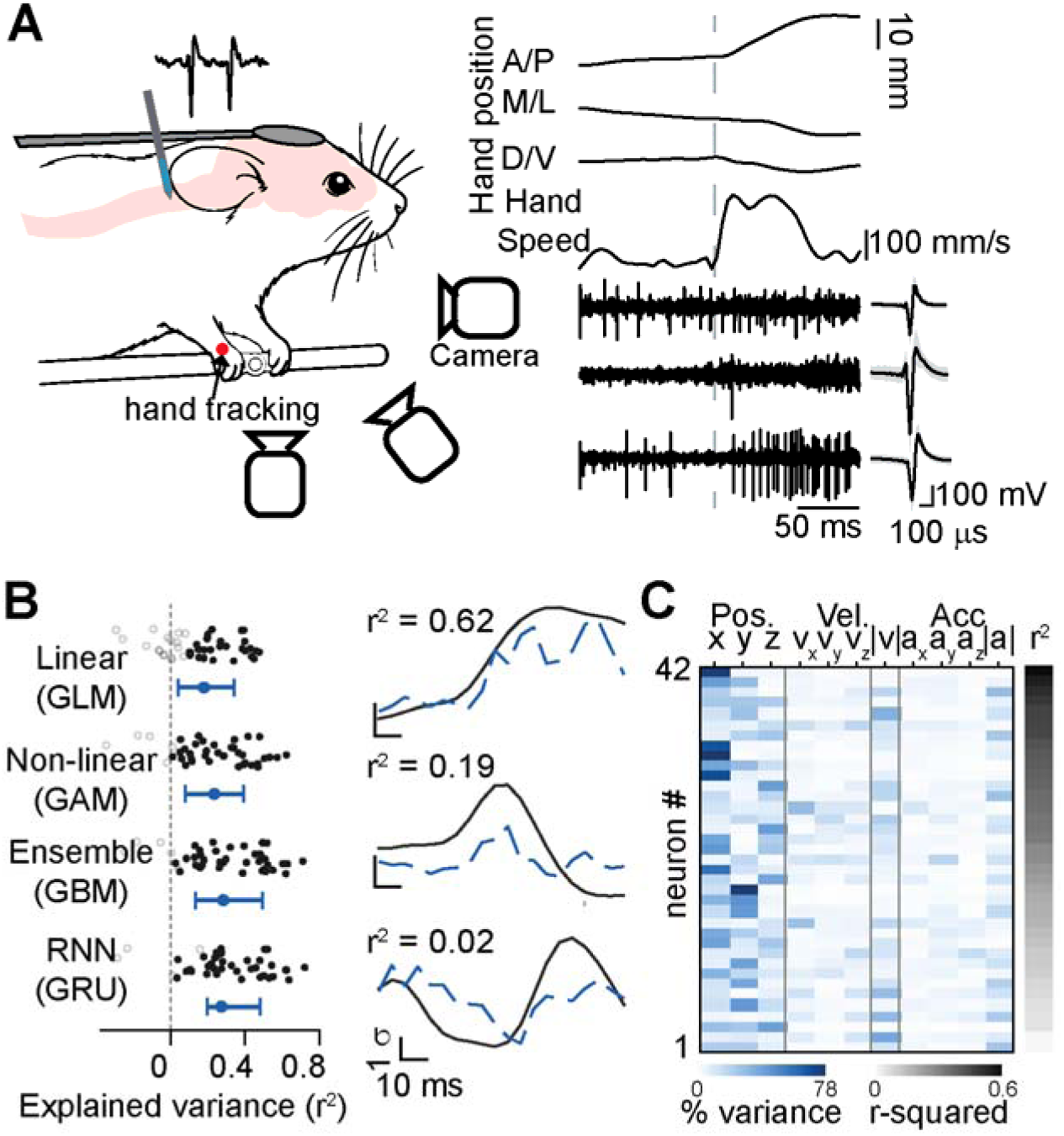
Cervical spinal cord contains neural representations of hand kinematics. **(A)** Left, Single-unit electrophysiology from medial deep dorsal horn of the cervical spinal cord in awake mice making spontaneous, un-instructed forelimb movements. Right, three example neurons aligned to 3d hand kinematics. **(B)** Left, Linear and non-linear encoding models show that most neurons (37/42, 2 animals) contain significant information about hand kinematics. Error bars are median +/- IQR across all recorded neurons. Right, example model fits of an interpretable non-linear model (Generalized Additive Model, GAM) model (dashed line) overlaid with smoothed z-scored firing rates for three example neurons. Neurons that carry significant information of hand kinematics in their firing rates are shaded black, the rest are shaded gray. Complex models can extract more information from neurons than simple models. **(C)** An interpretable non-linear model (GAM) shows that hand position is largest component of the information in the neural firing rates. Neurons are ordered by how much variance in their firing rates can be explained by hand kinematics (gray-black). The contribution of each kinematic parameter to the overall explained variance of the neuron is represented by the shading (white-blue). x, y, z are the A/P, D/V and M/L position of the hand relative to the limb’s position on the perch. Neurons were considered to have significant kinematic information if their r-squared values were both significantly above 0 and higher than values a circularly shifted bootstrap (p<1e-6) (Methods).

Recordings in anesthetized mice during passive movement provide a way to determine the sensory code directly and establish a template onto which other signals, such as efference copy and descending commands, can then be layered. To this end, we developed a motion control system that precisely moved the forelimb of an anesthetized mouse with micron-millisecond resolution (Fig. 2A, left and Fig. S2A-D) while systematically recording stable high-density, single unit electrophysiological signals (Fig. S2E-G). Neurons were sampled at different mediolateral and dorsoventral locations of the ipsilateral cervical spinal cord (Fig. 2B, left). Neurons encoding kinematic information were predominantly localized to medial lamina IV and V (Fig. 2B, right), where more than half the neurons were modulated by movement (53%, 145/273), and medial lamina VII/lateral lamina VIII, a region known to contain premotor interneurons that receive muscle and tendon afferent input(*19*), where approximately a quarter of the recorded neurons responded to movement stimuli (19/80). The patterns of neural activity represented by movement-modulated neurons were then examined using two complementary motion stimuli patterns that emphasized either position or velocity, two major kinematic parameters that contribute to proprioception, in a larger neural population (1570 neurons, 41 mice). First, we densely sampled the movement space by moving the limb in 64 directions at 40 mm/s (Fig. 1A, middle). We then sampled the velocity space by moving the limb in 8 directions at 20 mm/s, 40 mm/s and 80 mm/s (Fig. 1A, right).

**Fig. 2.**
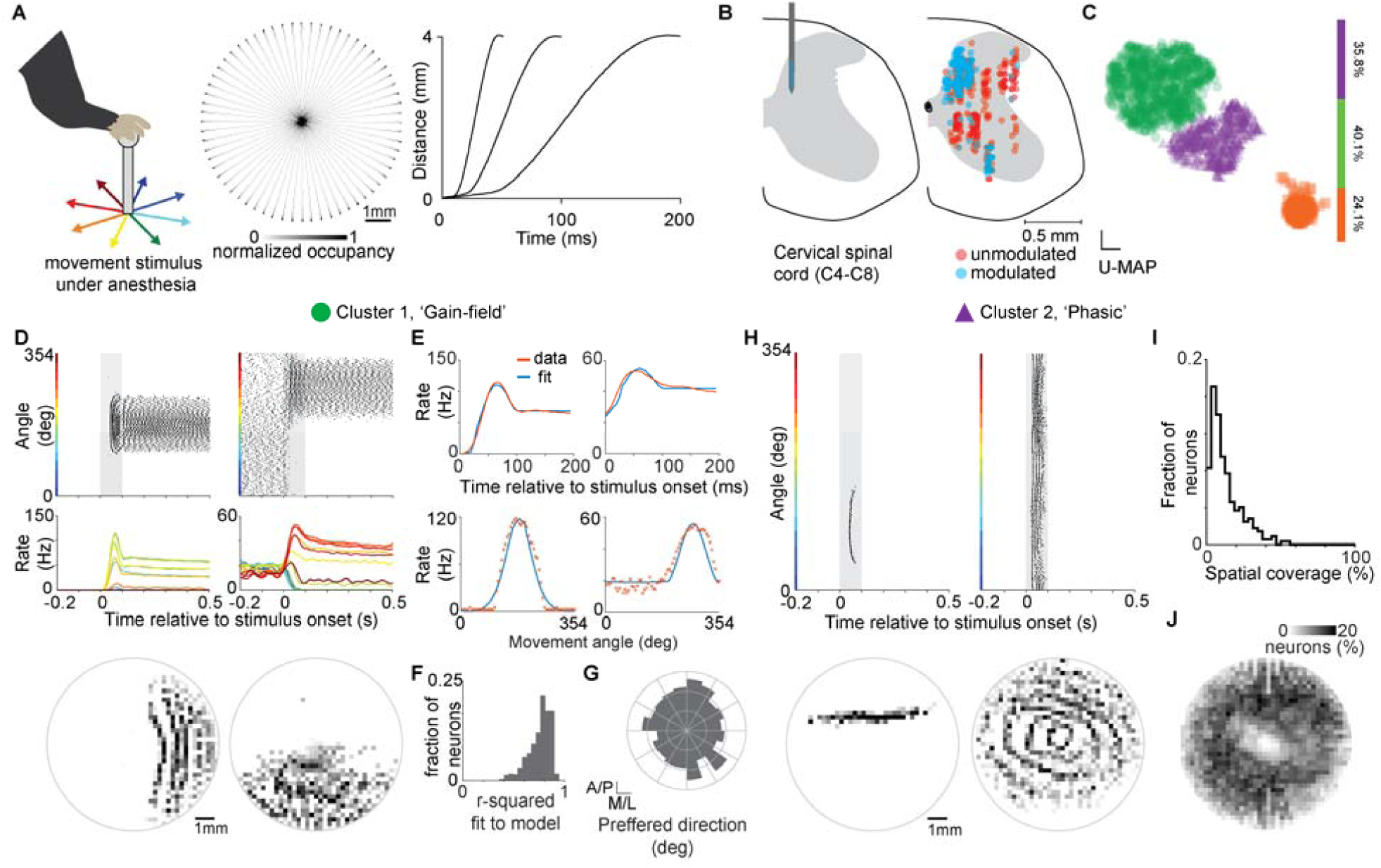
Electrophysiology in anesthetized mice reveals precise sensory representations of forelimb kinematics. **(A)** Left, The forelimb of an anesthetized mouse was systematically moved by a precise motion control system (Fig. S1) to densely sample the spatial and temporal dynamics of movement. Middle, spatial histogram of 1,920 trials recorded from a single session. Right, the three minimum-jerk speed profiles used in the motion stimuli. **(B)** Left, a schematic for *in vivo* multielectrode array recordings of single unit neural activity from the cervical spinal cords of mice. Middle, neurons modulated by the motion stimulus were localized to the medial deep dorsal horn and the lamina VIII of the cervical spinal cord. (n = 524, 14 mice) **(C)** Unsupervised clustering of the peristimulus time histogram of modulated neurons (n = 1570, 41 mice) reveals three distinct clusters: Gain-field, Phasic and Cutaneous-responsive neurons. **(D)** Example raster, rate histogram and spatial firing fields for two example Gain-field neurons. Trials sorted and grouped by angle of movement stimulus. 10 trials per movement direction. Shaded region indicates duration of movement stimulus. **(E)** Temporal (Top) and directional (Bottom) fits of the rate histograms of example neurons in (d) to a gain-field model for position, speed and direction of motion stimulus. **(F)** r-squared values for fits of Gain-field neuron rate histograms to a gain-field model. **(G)** Preferred response directions of the Gain-field neurons are widely distributed. **(H)** Example raster, rate histogram and spatial firing fields for two example Phasic neurons. Trials grouped as in (D). **(I)** The sizes of the spatial firing fields of Phasic neurons have a fat-tailed distribution. **(J)** Spatially tiled responses of Phasic neurons have complete coverage of the stimulus space.

By performing unsupervised clustering of the neural responses (Fig. 1C, Methods), we found three major clusters. The firing rates of Cluster 1 neurons contained a pronounced peak at the neuron’s preferred reach direction and smooth decreases as the movement angle deviated from this value (Fig. 2D, Fig. S3A). During movement, their responses showed a dynamic component that transitioned into a slowly adapting static component at the end of the reach as the hand was held at a static position (Fig. 2C). Changing the speed of the movement stimulus altered the dynamic component, but not the final static component (Fig. S3A). These observations showed that the firing rates of neurons in Cluster 1 were jointly modulated by the direction, speed and position of the forelimb (Fig. 2D, Fig. S3A). A gain-field rate model with a von Mises function for directional tuning that accounted for differential speed and position components explained >90% of the variance in firing rates of individual neurons (Fig. 2E-F, Fig. S3B, Methods). We therefore term these units ‘Gain-field’ neurons. While some Gain-field neurons had a large dynamic (speed related) component, others were more sensitive to the static (position related) component (Fig. S3C). Across the whole population, the speed/position component ratio was a single peaked Gaussian distribution (Fig. S3D, Methods). Most (∼84%) Gain-field neurons were tuned to a single direction (Fig. 2D, Fig. S3E) and, as a population, the distribution of tuning directions was largely uniform (Fig. 2G). A minority were tuned to two directions (∼14%) and very few (<2%) to more than two directions (Fig. S3C,E). The width of the tuning directions for these neurons were relatively broad, with a median dispersion of ∼50 degrees (Fig. S3F).

Cluster 2 neurons had firing fields at precise spatial locations, with some neurons firing a single spike with striking precision (∼5μm) across all trials (Fig. 2H left, Fig. S4A). This cluster also contained a subpopulation of the neurons that fired in precise radially-periodic firing fields over the entire workspace (Fig. 2H right, Fig. S4E). Neurons in this cluster lacked any baseline activity and typically had firing fields restricted to <10% of the workspace (Fig. 2H,I). We term these neurons ‘Phasic’. Most importantly, some Phasic neurons had precise representations that occupied varying spatial scales (Fig. 2I, Fig. S4A-E), i.e. ‘Multiscale representations’ - a key feature for a basis-set that is computationally efficient and scale invariant(*20*). To determine if the Phasic neurons spatially tiled the kinematic workspace, we estimated their firing fields (Methods) and overlaid them on a single grid (Fig. 2J). This analysis showed that their composite neural activity provided good coverage of the spatial workspace, with each location represented by ∼10% of the Phasic neurons.

In contrast to Clusters 1 and 2, neurons in Cluster 3 exhibited relatively noisy and inconsistent responses to the movement stimulus (Fig. S5), and their firing rates were only weakly tuned to kinematics. We hypothesized that these neurons were responding to stimulation of skin and hair induced by the motion of the arm, a prediction that we subsequently tested and confirmed. We putatively term these ‘Cutaneous-responsive’ neurons.

To rule out the possibility that these motion-specific neural responses were inherited from supraspinal structures, we recorded responses to the movement stimulus after making complete spinal transections at the C4 cervical segment that physically disconnected the brain from the spinal cord (Fig. S6A). Importantly, we saw abundant numbers of both Gain-field (56/110, 2 mice) and Phasic neurons (43/110, 2 mice) in the cervical cords of mice following the complete transection of the spinal cord (Fig. S5B,C), thereby demonstrating that these representations of forelimb movement in the spinal cord are constructed locally and are not inherited from the brain.

### Gain-field and phasic neurons are molecularly distinct

The distinct features of Gain-field and Phasic neurons suggested that they are likely to be different cell-types. To resolve these cell-types, we used optogenetic-assisted identification (Fig. 3A) of four molecularly-defined interneuron (IN) populations (Lbx1 (dI4-dI6 INs), Ptf1a (dI5 INs), PV (dPV INs) and Calb1 INs) that reside in the dorsomedial region of the spinal cord and are proposed to receive proprioceptive information. *Lbx1*+ INs spanned all three classes (Fig. 3B,C), *Ptf1a*+ neurons were Phasic or Cutaneous but lacked Gain-field responses (Fig. 3D,E). Both *PV*+ (36/40) and *Calb1+* (39/40) neurons were largely unmodulated. *Lbx1+/Ptf1-*Gain-field and *Lbx1+/Ptf1a+* Phasic neurons are therefore molecularly- and developmentally-distinct populations of dorsally-derived sensory interneurons

**Fig. 3.**
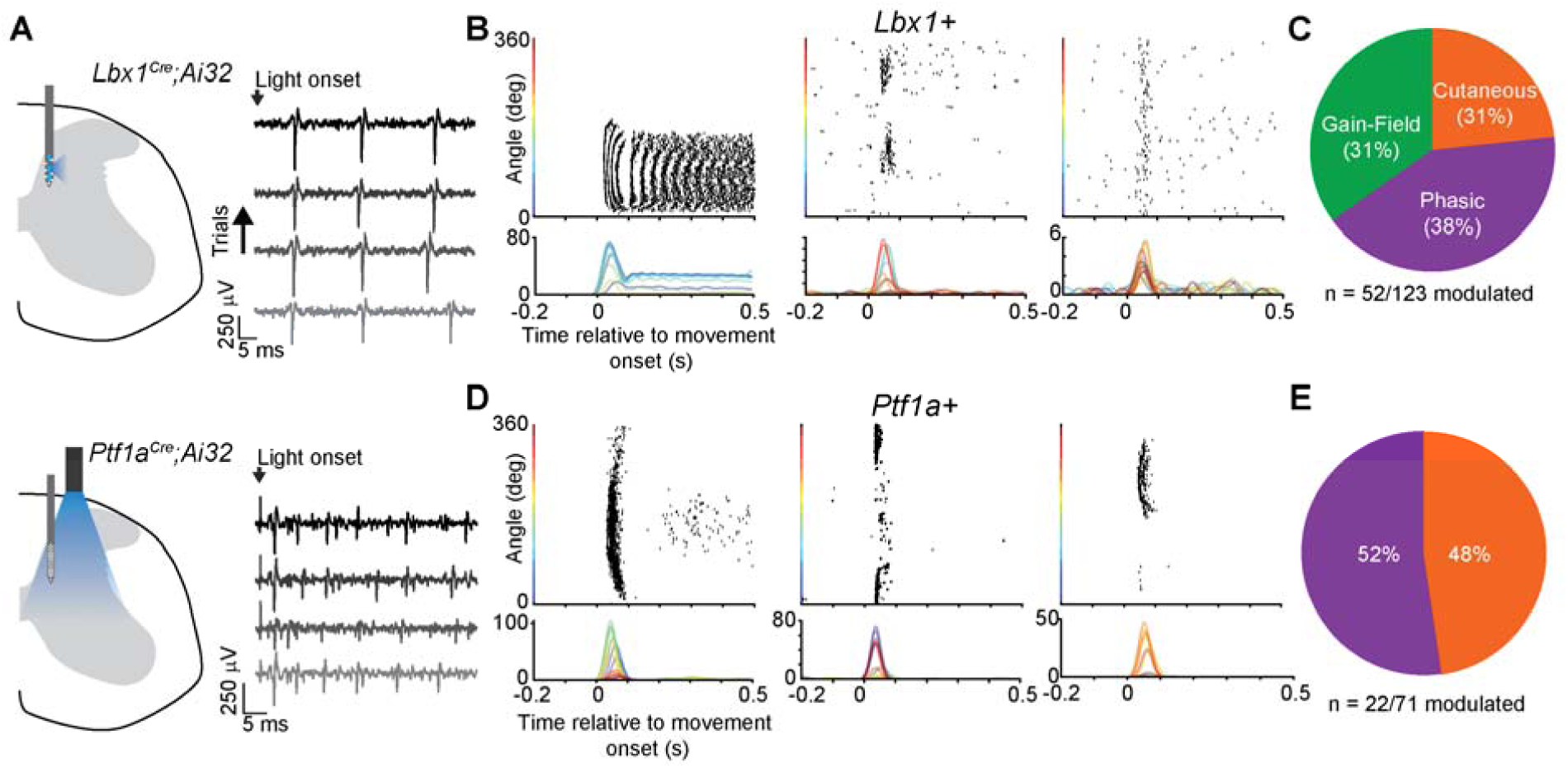
Gain-field neurons and Phasic neurons are molecularly distinct. **(A)** Left, Optogenetics-assisted identification using localized and surface stimulation for excitatory and inhibitory neural populations, respectively. Right, examples of light-responsive neurons across four trials. **(B-C)** Lbx1+ neurons encompassed all three neural response classes and in approximately the same proportions as the general populationv. (B) Example raster of Lbx1+ Gain-field (left), Phasic (middle) and a Cutaneous-reponsive (right) neuron. **(D-E)** Ptf1a+ neurons were restricted to Phasic and Cutaneous-reponsive neurons. (B) Example raster of two Phasic-neurons (left. middle) and a Cutaneous-reponsive (right) neuron.

### Identification of a neural geometry for forelimb proprioception

To understand how the neural population activity evolves over the trajectory of the movement and the relationship between the population response to movements in different directions, we used de-mixed principal component analysis(*21*) (dPCA) on the smoothed firing rates of neurons where both the velocity and movement space were densely sampled (n = 431 neurons, Methods). dPCA is a dimensionality reduction method that can separate the population neural activity into components associated with specific variables, such as direction. 72% of the neural variance could be described by 4 dimensions (Fig. S7A). Although the dPCA algorithm did not have access to the values for the direction, speed or position of the movements, the neural population structure was an ordered representation in this low dimensional space (Fig. 4A) with a pinwheel structure that faithfully reflected the kinematic state of the forelimb (Supplemental Video 2). Specifically, we found that the first three condition dependent components correspond to the anterior/posterior position, medial/lateral position and the speed of the hand movements, respectively (Fig. 4B). Importantly, this order was maintained in movements with different peak speeds (Fig. S7B,C).

**Fig. 4.**
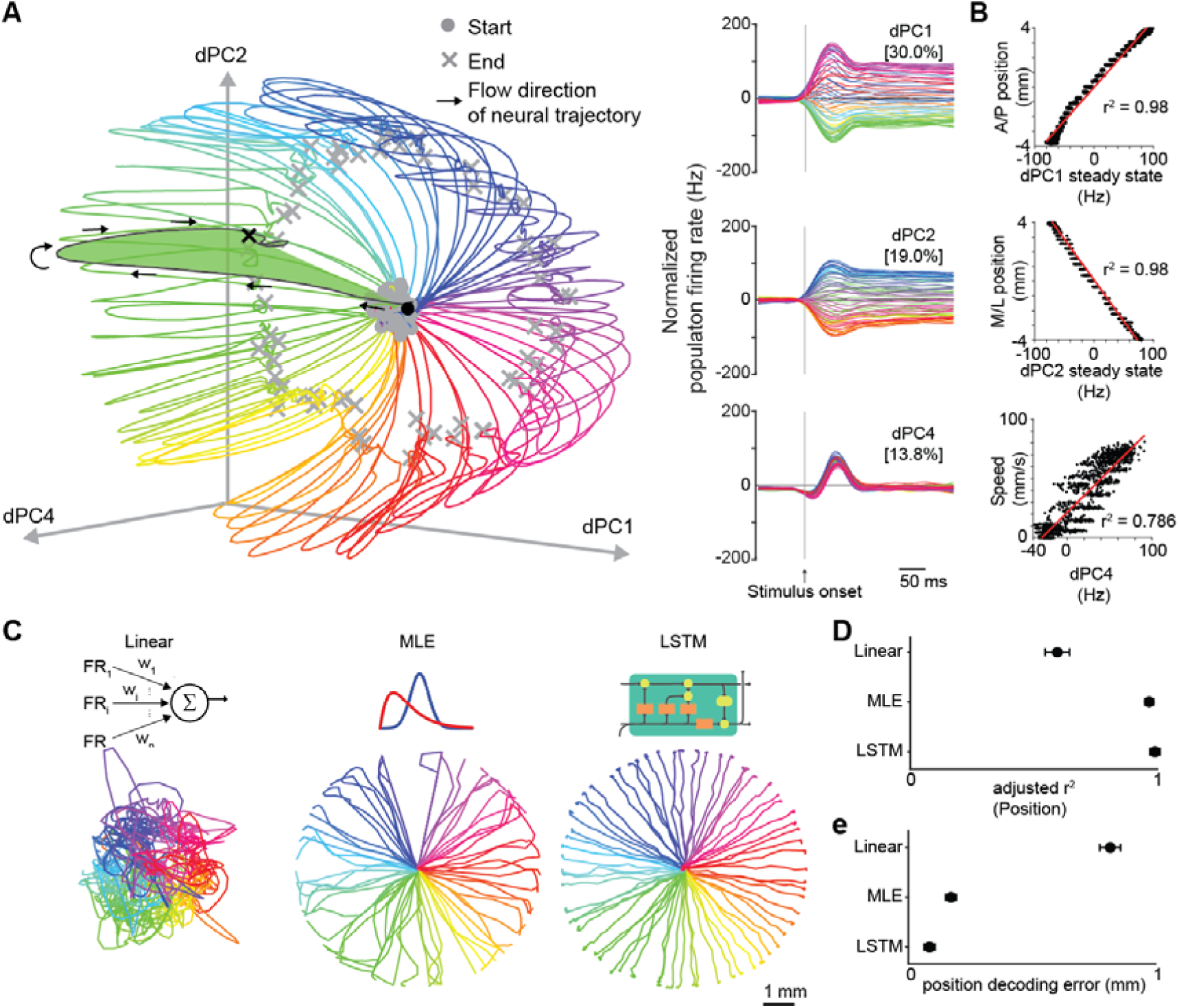
Neural population code in the cervical spinal cord is an ordered representation of forelimb kinematics. **(A)** Left, three condition-independent demixed principal components (dPCs) of neural activity in the cervical spinal cord color coded by the direction of stimulus. Right, temporal dynamics of the same three dPCs. **(B)** The dPCs are strongly correlated to specific kinematic variables, such as anterior-posterior position (dPC1), medio-lateral position (dPC2) and speed (dPC4). **(C)** Example decoding of single trial kinematics from neural recordings in the cervical spinal cord using Linear (left), maximum-likelihood (MLE)(middle) and a long-short term memory (LSTM) recurrent artificial neural network (Right) models. **(D) and (E)** r-squared values (D) and the mean absolute error (E) for single trial decoding across the three model classes. Values are represented as Mean +/- SEM.

We then evaluated the relative contribution of each of the three identified neural clusters to the construction of this low dimensional manifold by performing dPCA on the individual clusters. This analysis showed that Gain-field and Phasic neurons make distinct contributions to the neural geometry, while the Cutaneous-responsive neurons have almost no ordered structure (Fig. S6D-F). These analyses demonstrate that this manifold does not arise trivially from the activity of any movement-modulated neurons (such as the Cutaneous-responsive neurons), but instead requires the patterns of activity exhibited by Gain-field and Phasic neurons.

The structure of the neural population response we discovered argues that the spinal cord faithfully represents the kinematic state of the limb. To further evaluate this possibility and identify the neural computations that are required to estimate the kinematics of the limb, we built and validated decoders to extract the forelimb kinematic state from the neural population activity (Methods). Three classes of decoders that approximate biologically plausible computational mechanisms (Fig. 4C, Fig. S8, Methods) were analyzed. Linear models that approximate feedforward neural circuits that compute using instantaneous firing rates of inputs(*22*), maximum-likelihood models (MLE) models which approximate associative networks that rely on probabilistic priors(*23*), and long-short term memory models (LSTM) that resemble recurrent neural networks(*24*) with the ability to incorporate temporal context. A compact LSTM network (8 hidden units) with access to information from ∼40 neurons was sufficient to decode single trial kinematics with an accuracy of ∼50 um for position and ∼5 mm/s for speed (Fig. 4D,E, Fig. S8E). In contrast, both the linear and MLE models performed significantly worse on single trials (Fig. 4D,E, Fig. S8A-D). These analyses demonstrate that downstream neural circuits with compact memory and recurrent connectivity can compute forelimb kinematics with high precision from the neural population activity we recorded in the cervical spinal cord.

### The neural code for kinematic state requires activity of muscle and tendon afferents but not cutaneous afferents

The spinal cord receives two major sources of mechanosensory information that have been implicated in estimating the kinematic state of the limb, mechanoreceptors embedded in muscles and tendons, and skin-derived cutaneous low-threshold mechanoreceptor (LTMR) afferents(*25*). To identify which of these sensory pathways are required to construct the kinematic neural code we report here, we employed an intersectional genetic approach to selectively target and ablate them. Muscle and tendon afferents were targeted with *Pirt::Cre*;*PV^FlpO^* (*26–28*) and cutaneous LTMRs with *Avil^CreERT2^;MafA^FlpO^*(*29*, *30*). Crossing to a *R26^ds-tdT^*reporter confirmed ≥90% of *Pirt::Cre*;*PV^FlpO^* afferents co-expressed PV and TrkC, a marker of muscle/tendon afferents (Fig. S9 A-D). *Avil^CreERT2^;MafA^FlpO^*labeled ∼19% of DRG neurons (Fig. S9F) which were large-diameter, non-nociceptive, and excluded muscle/tendon afferents. MafA marks Aβ LTMRs including rapidly adapting Meissner- and Merkel-associated afferents(*29*), and is not expressed in Aδ-LTMRs or in unmyelinated C-LTMRs (*31*). Aβ afferents are the primary cutaneous afferents implicated in proprioception in humans and cats(*32–35*).

We then selectively ablated either the cutaneous, or the muscle and tendon afferents using the *Tau^ds-DTR^* system(*36*) while recording motion stimulus driven neural activity in the cervical spinal cord (Fig. 5A-F, Methods). Ablating muscle and tendon afferents led to the complete absence of neurons displaying phasic responses and a near absence of neurons displaying gain-field responses (Fig. 5H, right). Cutaneous-responsive neurons with low kinematic information formed the vast majority of residual neural responses. Conversely, the proportion of Cutaneous-responsive neurons were more than halved in mice lacking most of their LTMR afferents while the Gain-field and Phasic neurons were retained (Fig. 5I, right). Consistent with these changes in afferent input, the neural geometry for proprioception was entirely disrupted by muscle and tendon afferent ablations (Fig. 5H, left), but was largely unaltered by ablation of LTMR afferents (Fig. 5I, left). These experiments demonstrate that muscle and tendon afferents provide the overwhelming majority of the kinematic state information of the limb, with cutaneous afferents making little, if any, contribution to the construction of the neural geometry for forelimb kinematics.

**Fig. 5.**
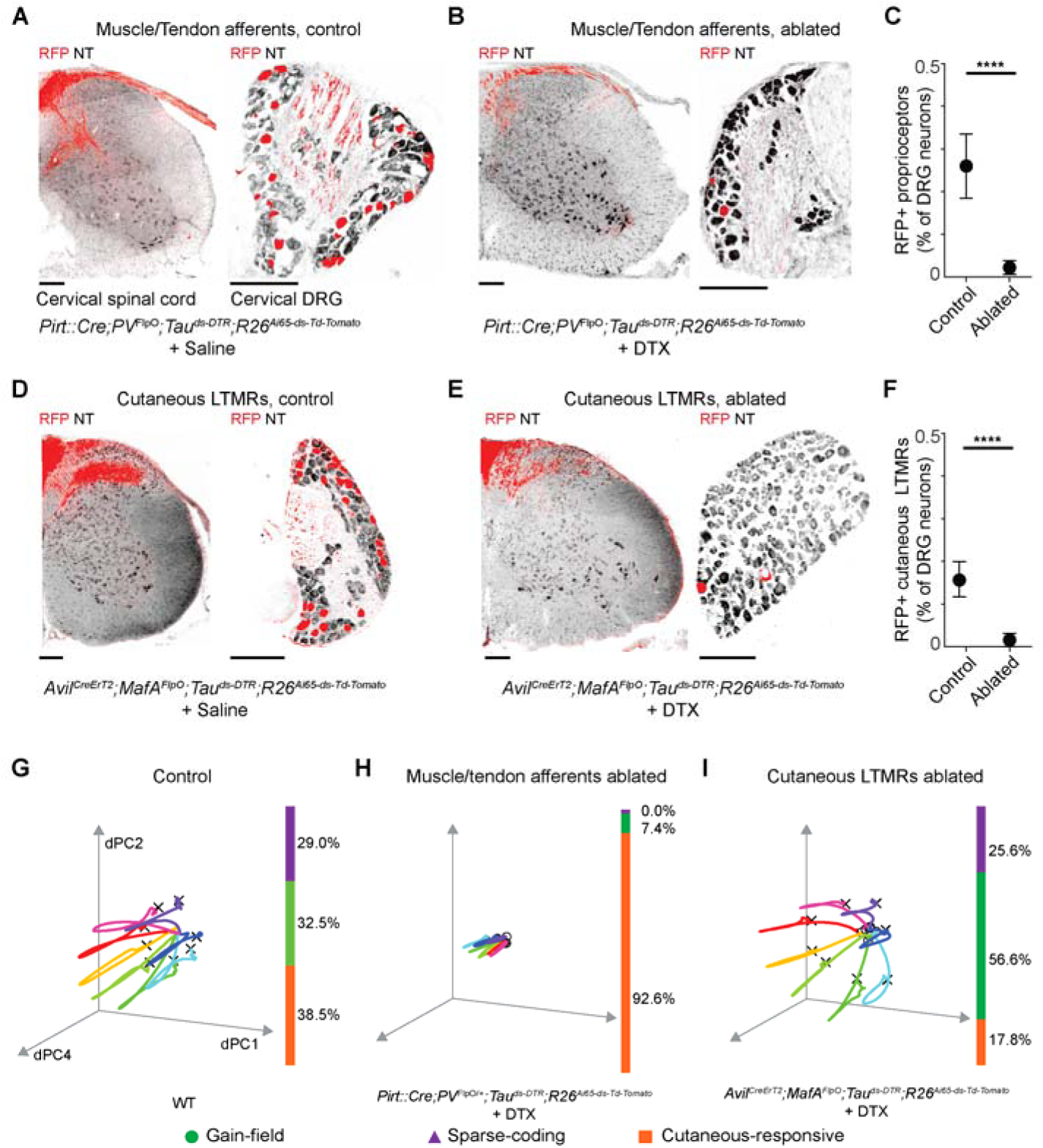
Neural code for forelimb proprioception requires muscle afferents, but not cutaneous afferents. **(A-C).** Selective ablation of muscle afferents by an intersectional genetic strategy with a cross of *Pirt::Cre, PV^Flpo^, Tau^ds-DTR^* and *R26^ds-tdT^* alleles. Representative images of cervical spinal cord and dorsal root ganglia with an infusion of saline (A) or diphtheria toxin (B). Quantification of the ablation efficiency across mice (D). **(D-F).** Selective ablation of a subset of cutaneous low threshold mechanosensory afferents by an intersectional genetic strategy with a cross of *Avil^CreERT2^, MafA^FlpO^, Tau^ds-DTR^* and *R26^ds-tdT^*alleles. Representative images of cervical spinal cord and dorsal root ganglia with an infusion of saline (D) or diphtheria toxin (E). Quantification of the ablation efficiency across mice (F). **(G-I).** Ablation of muscle/tendon afferents and cutaneous LTMR afferents target distinct clusters of neural responses and differentially perturbs the neural geometry. (H) Ablation of muscle afferents collapses the neural geometry for proprioception (left) by specifically eliminating gain-field and phasic neural responses (right) compared to controls (G), whereas (I) ablation of cutaneous LTMRs only affects Cutaneous-responsive neurons but not gain-field or phasic neural responses, leaving the neural geometry for proprioception intact. Color bars indicate percentage of the gain-field, cutaneous and phasic neurons observed in the recorded population. Values in (C),(F) are represented as Mean+/-SD. **** denotes a p-value <0.0001 for a two-sided t-test on the means. n = 10 sections from at least 3 animals each.

### Reducing muscle and tendon afferent input perturbs end-point precision but not reach direction

The control of voluntary movement is hypothesized to rely on three essential signals: feedforward commands, efference copies and sensory feedback(*37*). Having identified the code for sensory feedback in the cervical spinal cord and its dependence on muscle/spindle afferents, we next asked how this sensory information is used during voluntary skilled movement. We reasoned that perturbations of sensory information would enable us to identify the specific contributions of sensory feedback to voluntary movement that cannot be compensated by feedforward motor commands or their efference copies. To this end, we trained freely moving mice to use a joystick and make rapid directional reaches to spatial targets(*38*) (Fig. 6A, Methods). These trained mice made highly stereotyped reaching movements, with precise end-points. Ablating muscle and tendon afferents led to a progressive degradation in the reaching movements over the following three days, characterized by reduced speeds and increased variability in both reach direction and end-point precision (Fig. S10A-D). By day three, muscle and tendon afferent-ablated mice were unable to initiate reaching and made almost no movements (Fig. S10B). In contrast, the reaching behavior with LTMR ablations (Fig. S10E-H) and in controls (Fig. S10I-L) was unaffected by DT administration. These data, in addition to previous reports(*39*, *40*), demonstrate that sensory feedback from muscles and tendons is essential for both initiating and coordinating skilled movement.

**Fig. 6.**
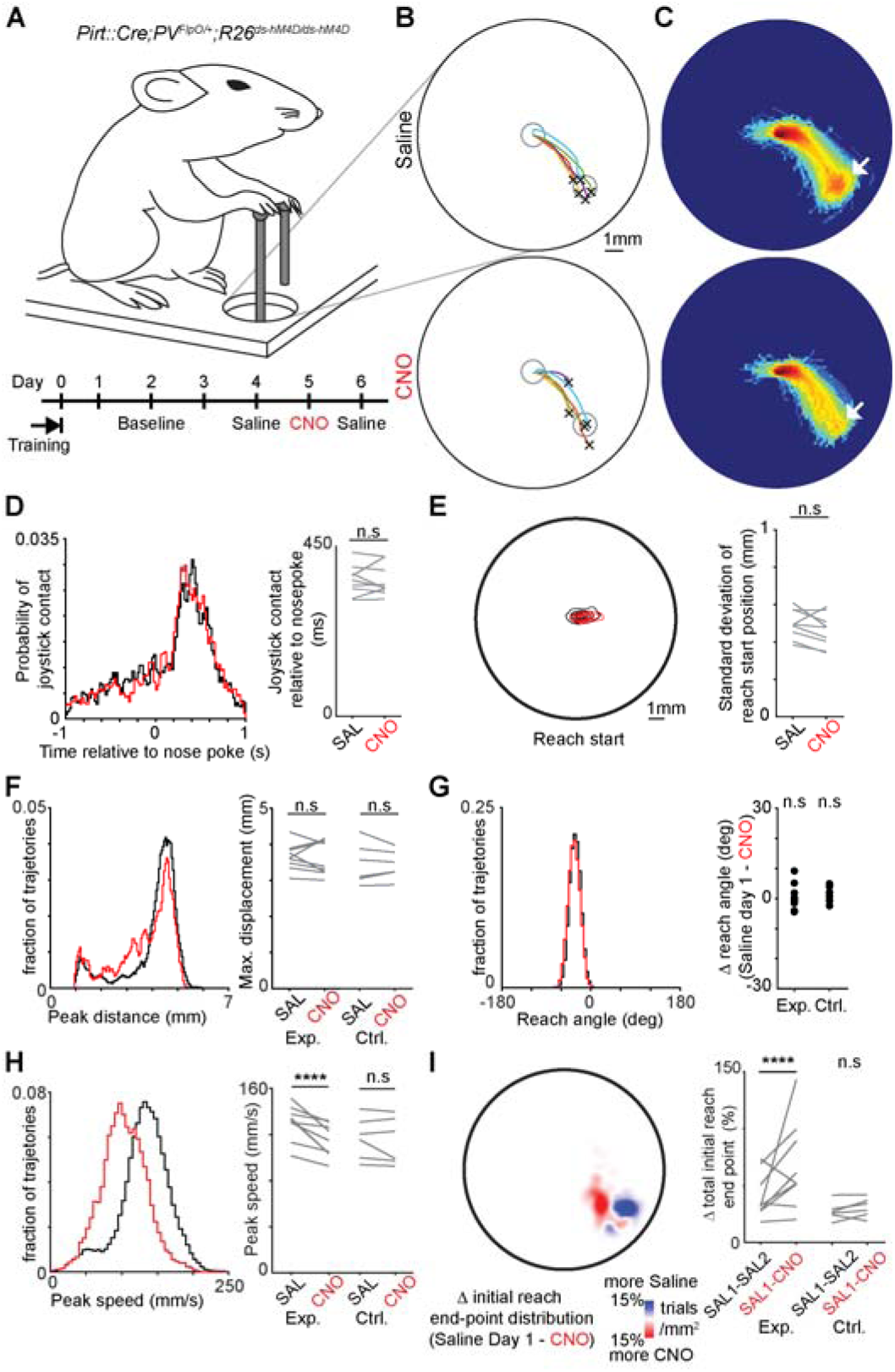
Transient silencing of muscle and tendon afferents perturbs speed and end-point precision, but not direction of skilled forelimb movements. **(A)** Freely moving mice expressing the hM4D DREADD in the muscle/tendon afferents were trained to make rapid center-out reaches to spatial targets using a custom-built low torque joystick (Top). After training, mice were given intraperitoneal injections of either saline or clozapine-N-oxide (CNO). **(B)** 5 example trajectories from saline (top) and CNO (bottom) sessions. The grey circles represent the starting position and target. X is the end-point of the reach. **(C)** Log-scaled heat maps for all the trajectories from a saline session (top) and a CNO (bottom) session. Note the reduced of density at the target for the CNO session (arrows). **(D-I)** Effect of CNO induced silencing on joystick movements. Time to joystick contact after nosepoke (D), distribution of joystick starting points (E), peak distance (F), reach angle (G), peak speed (H) and (I) end-point distributions. Data from an example experimental animal (left) and across all animals (right). Experimental animals (Exp.) had both the Cre and FlpO alleles, Control animals (Ctrl.) had either the Cre or the FlpO allele. **** denotes a p-value <0.0001 for a hierarchical bootstrap test on the medians, n = 9 and 7 animals for the experimental and the control cohort, respectively.

Given the limited insight these experiments provided regarding the use of information from muscle afferents in the control of skilled movement, we developed a subtler perturbation using DREADD-based chemogenetics. The intersection of *Pirt::Cre, PV^FlpO^* and *R26^ds-tdT-hM4D/ds-tdT-hM4D^*was used to selectively express the Gi-coupled hM4D receptor in muscle and tendon afferents (Fig. S11A). We validated the effectiveness of the DREADD receptor electrophysiologically by recording motion stimulus-evoked neural activity in the cervical spinal cord from the same neurons prior to, and after, CNO-treatment (Fig. S11 B,C). CNO infusion reduced firing rates in the Gain-field neurons by ∼30% on average (Fig. S11 B,D). While the firing rates were reduced, the Gain-field neurons maintained their directional tuning (Fig. S11 B). This reduction in firing rates of Gain-field neurons alters the position and speed components of the neural code, but not the direction. Phasic neuron responses deteriorated more than those of Gain-field neurons, and this was accompanied by a marked disruption of their firing fields (Fig. S11 C,E), thereby removing precise positional feedback during movement. The combined effect of these disruptions to the neural code across Gain-field and Phasic neurons was a loss of precise speed and positional feedback during movement, but not directional information. If neural circuits controlling voluntary movements rely on these kinematic estimates for precision, then we posited that sensory perturbations during reaching would impair components that require precise positional feedback such as end-point accuracy, while leaving the overall reach direction intact. To test this prediction, we trained *Pirt::Cre; PV^FlpO^; R26^ds-tdT-hM4D/ds-tdT-hM4D^* animals in the joystick reaching task and transiently silenced their muscle and tendon afferents. In control sessions with saline infusion, mice made highly stereotyped movements. They nose-poked, grasped the joystick with the right forelimb (Fig. 6A,D,E), and made a series of rapid, accurate reaches to targets (Fig. 6B,C, top). Activation of hM4D by CNO infusion did not impair the stereotypy of joystick grasp time relative to nosepoke (Fig. 6D) or change the variability of the reach initiation point (Fig. 6E). In contrast to the ablation experiments, mice were therefore able to contact the joystick equally well with hM4D activation. As predicted by the deficits in the neural code, hM4D activation altered the trajectories of reaching movements (Fig. 6B,C, bottom). Overall reach distance and direction were not significantly altered (Fig. 6F,G), but reaches were slower (Fig. 6H) and showed substantial errors in endpoint (Fig. 6I). These deficits reversed in the following saline session. Mice lacking the hM4D allele showed no deficits after CNO (Fig. 6F-I), arguing against off-target effects of CNO. This selective disruption of endpoint accuracy, with distance and direction unchanged, is consistent with a key behavioral prediction of the neural code we have defined.

## Discussion

Our study shows that neurons in the cervical spinal cord generate an ordered neural population code with a pinwheel structure that encodes a precise moment-by-moment estimate for kinematic state of the forelimb. This representation is constructed by two classes of neurons with distinct firing patterns. It comprises of a gain-field rate code that encodes speed, direction and position and a Phasic code that encodes specific spatial locations. Genetically targeted manipulations of sensory inputs show that muscle and tendon afferents, but not cutaneous afferents, are required to construct this neural code. Moreover, transient perturbations of muscle and tendon afferents perturb end-point accuracy in skilled movements. Our findings show that the spinal cord uses sensory information from muscle and tendon afferents to perform the complex computations required to estimate the kinematic state of the forelimb, which is required for skilled movement.

Until now, stable recordings from the cervical spinal cord of awake behaving animals have been difficult to obtain. Moreover, they have not been possible in mice as the cervical cord is difficult to access, and recordings are corrupted by artifacts induced by breathing and limb movement. Our spinal implant overcomes these limitations by providing chronic access to the cord and stabilizing its motion relative to the recording probe, which enabled stable, simultaneous recordings from populations of neurons during forelimb movement. Even so, awake recordings are confounded by additional signals layered onto the proprioceptive code. These include descending command signals, efference copies, other sensory modalities and behavioral state information. This was evident in our encoding analysis where even flexible nonlinear models explained only a modest share of firing-rate variance in awake animals. Cross-validated R^2^ reached about 0.75 in the best-fit neurons but fell below 0.25 in most neurons (Fig. 1), indicating they are integrating kinematic signals with other inputs. Importantly, the Gain-field and Phasic neurons we characterized under anesthesia were also present in awake animals (Fig. S1C) and showed similar tuning properties, indicating that the coding properties we describe reflect the intrinsic organization of the spinal circuit that is maintained under anesthesia. The use of anesthesia therefore allowed us to isolate the underlying sensory code of each neuron class, characterize its properties in detail, and establish a ground truth for understanding how other signals could layer onto and modulate it in the behaving animal.

The firing patterns we see for Gain-field neurons in the ‘preferred direction’ are similar to those observed in isolated muscle spindle Ia and II afferents in response to stretch(*41*, *42*), indicating that Gain-field neurons maintain the temporal precision of the information transmitted from the muscle afferents. However, contrary to predictions made by computational models(*43*) and prior observations in the lumbar spinal cord(*44*), the directional tuning of these neurons is largely uniform for the entire workspace and is not strongly biased along one or two axes. A uniform representation of reaching directions has previously been interpreted as a signature of higher-order computation(*43*). Somewhat surprisingly, we didn’t see distinct clusters of firing rate patterns that would correspond to Ia, II and Ib afferents. Rather, the position/speed ratio forms a Gaussian distribution, suggesting that Gain-field neurons integrate proprioceptive information from all three afferent types before transmitting it to downstream structures. This is consistent with studies in the cat showing that many spinal interneurons receive convergent Group Ia, Ib and II afferent input(*45*, *46*), and the integration of sensory signals in the ventral nerve cords of flies(*47*). However, a direct test of whether gain-field neurons integrate across afferent types will require selective perturbation of each type, this will require the development of genetic tools based on recently identified molecular markers of Ia, Ib, and II afferents (*48*, *49*). Intriguingly, the firing patterns of these spinally computed Gain-field neurons are similar to those seen in the primate S1 cortex(*7*), suggesting that kinematic representations in higher-order structures are inherited from the spinal cord. The Phasic neurons we have identified constitute an entirely novel class of neural responses for proprioception. These neurons tile the entire workspace and are important for positional precision during movement, with the firing of a subset of these neurons able to immediately clarify a limb’s spatial location.

The firing patterns of Gain-field neurons are consistent with a rate code, with their persistent activity during and after moving the forelimb (Fig. 1) being well suited for generating a neural population state that represents a specific movement context, e.g. posture. On the other hand, the information embedded in the Phasic neurons is encoded near-instantaneously by the precise timing of a single spike(*50*), which is consistent with a spike timing code. This however makes information from Phasic neurons sensitive to noise and transient with no memory. Rate codes and spike-timing codes have different properties and utility for information processing(*51*). Rate codes are noise robust(*52*) and are able to encode graded information by modifying firing rates, however they require integration over longer times scales (∼10 ms) or across many neurons. We propose that the presence of these two complementary coding mechanisms in the spinal cord enables neural circuits in the central nervous system to leverage the benefits of both. The results of our behavioral experiments provide direct evidence for this. Chemogenetic silencing disrupted firing patterns in Phasic neurons while leaving the activity in Gain-field neurons largely intact. The behavioral consequence for this perturbation was a loss in end-point precision, without altering direction and distance. These results indicate that the temporal precision of Phasic neurons is essential for accurate movements, and they delineate the differential contributions of Gain-field and Phasic neurons to the kinematic code.

The sharp distinction in firing patterns between Gain-field and Phasic neurons suggests they are separate cell types, and our optogenetics-assisted identification supports this. *Lbx1*+ neurons were movement-responsive and spanned all three classes, whereas *Ptf1a*+ neurons were restricted to Phasic and Cutaneous-responsive units. Our marker analysis identifies *Ptf1a+* neurons as Phasic neurons and defines Gain-field neurons by exclusion (*Lbx1+/Ptf1a-*), demonstrating the neural coding of kinesthesia is defined by developmental lineage rather than cell position. It is notable that Phasic neurons fall within an inhibitory lineage, since in canonical sparse-coding circuits, such as HVC projection neurons (*53*, *54*) and Kenyon cells (*55*, *56*), the sparse code is carried by excitatory neurons. Ptf1a-derived inhibitory neurons are also required for spinal sensorimotor adaptation in which the limb learns a spatial location (*57*), consistent with this lineage being necessary for learning limb position. Importantly, a number of dorsal neuron subtypes in the same area did not respond. *PV*+ dPVs, a glycinergic subpopulation of *Lbx1+* INs located in the medial deep dorsal horn, along with excitatory *Calb1*+ neurons in the same territory, were largely unmodulated by passive limb movement.

At the population level, the gain-field and phasic neurons together construct an ordered low-dimensional manifold. The pinwheel neural geometry in the cord likely reflects the geometry of the stimulus, and we would not expect it to persist for movements with a different geometry, such as a figure-eight. Its form was nonetheless not a foregone conclusion. Muscles differ greatly in size and spindle complement (*58*), and the mouse triceps is roughly four times the mass of the biceps (*59*). Modeling and recordings from muscle afferents indicate that the directional tuning of proprioceptive input follows muscle geometry (*43*), which predicts that some directions would be over-represented and would distort the population code. Instead, we observe a largely symmetric pinwheel (Fig. 4), consistent with the uniform distribution of direction tuning across single neurons (Fig. 2). This suggests that spinal circuits may contain intrinsic mechanisms that equalize the contribution of different directions, for example by integrating convergent afferent input into a representation of the limb rather than of individual muscles. Likewise, the alignment of dPC1, dPC2 and dPC4 with the AP position, ML position and speed of the limb was not imposed by the analysis, because dPCA has no knowledge of the stimulus coordinate frame. It indicates that the cord organizes its population activity along the axes of limb movement.

Prior modeling work on neural manifolds in motor control offer a complementary theoretical context for our results(*60*). It is known that low-dimensional manifolds can emerge when recurrent neural networks (RNNs) are trained on realistic muscle patterns or kinematics(*24*). These studies support our finding of an ordered spinal manifold, while also suggesting that manifold organization may be a general principle across the sensorimotor hierarchy, including the spinal cord. However, these modeling studies, while providing a normative framework for how structured low-dimensional representations can arise from task demands or network optimization, do not yet encompass the computational diversity, or the temporal precision exhibited by spinal Phasic neurons. Consequently, the combination of rate-based gain-field coding and Phasic observed here suggests that future models will need to incorporate parallel, complementary population codes to capture the richness of biological proprioceptive processing. The simultaneous presence of these two dichotomous information streams suggests that the downstream neural circuits can leverage the benefits of both, thus enabling the central nervous system to generate context-aware, rapid, goal-directed movements.

In assessing the contribution of cutaneous inputs to the kinematic code, it is notable that ablating MafA-defined Aβ LTMRs did not alter the code or skilled reaching (Fig. 5, Fig. S10E), as cutaneous afferents have been shown to carry kinematic information in cats(*34*, *35*), humans (*32*, *33*, *61*), and non-human primates (*62*). Two features of our study may explain the difference. First, the extent to which kinematic information carried by the afferents is transmitted to spinal interneurons in this species is not known. Second, cutaneous input appears to matter most at distal joints (such as fingers and wrist)(*62*), whereas muscle spindle information at proximal joints (Elbow and shoulder) is likely to be less ambiguous(*33*).

In comparing neural responses in medial lamina IV-V between awake and anesthetized mice, a key observation is that these regions are densely innervated not only by muscle and tendon afferents but also by descending inputs from motor and somatosensory cortex (*18*, *63*). This raises the possibility that the motion responsive neurons in these areas serve as neural substrates for top-down modulation by supraspinal structures, either for gain-modulation or suppressing reafference(*64–67*). Such inputs would facilitate the regulation of proprioceptive information in a task-dependent manner, particularly during dexterous movements that are rely on descending cortical control.

## Materials and Methods

### Mouse Lines

All experiments were carried out in accordance with NIH guidelines and were approved by the Salk Institute’s Institutional Animal Care and Use Committee. Mice were between 16-52 weeks of age (median 36 weeks) and housed in groups of up to 5 animals under a 12-h light/dark cycle till the day of the recordings. For behavioral experiments, mice were individually housed under a 12-h light/dark cycle for the duration of the study, and were tested during the dark phase.

All the mouse lines used in this study have been previously described, including Pirt::Cre, Avil^CreERT2^; PV^FlpO^, MafA^FlpO^, Ai65 (R26^FSF-LSL-tdTomato^, referred to here as R26^ds-tdT^), Tau ^FSF-LSL-DTR^ and R26^FSF-LSL-hM4Di^ (referred to here as R26^ds-tdT-hM4D^). The Cre and FlpO driver lines were maintained on a C57Bl/6 background, while the effector lines were maintained on a mixed background.

For the electrophysiology, we used the following numbers of mice per strain:

Wild type BL6(6), Pirt::Cre;PV^FlpO^;Ai65;R26^ds-tdT-hM4D^(14), Pirt::Cre;PV^FlpO^;Ai65;Tau^ds-DTR^(3), Avil^CreERT2^;MafA^FlpO^;Ai65;Tau^ds-DTR^(2), Lbx1^Cre^;Ai32 (4), PV^Cre^;Ai32 (3), Calb1^Cre^;Ai32 (3), Ptf1a^Cre^;Ai32 (6).

For behavior:

Pirt::Cre;PV^FlpO^;Ai65;Tau^ds-DTR^(4), Avil^CreERT2^;MafA^FlpO^;Ai65;Tau^ds-DTR^(4), PV^FlpO^;Ai65;Tau^ds-DTR^(2), Pirt::Cre;Ai65;Tau^ds-DTR^ (2), Pirt::Cre;PV^FlpO^;Ai65;R26^ds-tdT-hM4D^(9), PV^FlpO^;Ai65;R26^ds-tdT-hM4D^(3); Pirt::Cre;Ai65;R26^ds-tdT-hM4D^ (4).

### *In vivo* Cervical Spinal Cord Electrophysiology

Mice were deeply anaesthetized with isoflurane (4%). Fur was trimmed over the skull and the dorsal cervical area, and mice were placed in a stereotaxic frame (Kopf Instruments). A heating pad was used to prevent hypothermia. Isoflurane was delivered at 1-3% throughout surgery; this level was adjusted to maintain a constant surgical plane. Ophthalmic ointment was used to protect the eyes. Lidocaine diluted to 0.5% was injected subdermally along the incision line of the scalp and over the cervical spinal cord. The entire shaved area was disinfected by a two-stage betadine and alcohol scrub, repeated three times. The scalp was then removed with surgical scissors to expose the skull, which was thoroughly cleaned. The skull was then roughened with the scalpel to improve adhesion to the dental cement. A craniotomy was then made over visual cortex (−3.5 AP ± 3 ML), and a ground electrode soldered to a gold pin (A-M Systems) was then implanted into the craniotomy to a depth of 1mm. The ground pin and a headplate were secured to the skull with Metabond dental cement. After the Metabond hardened, a second incision was performed over the cervical vertebrae and the adherent fascia was gently removed from the underlying muscle. Muscle and connective tissue overlying the C4-C8 vertebrae were removed by blunt dissection until the vertebrae were seen. Tendons connecting to the dorsal surface of the vertebrae were disconnected and the facet joints were exposed. The vertebrae and the surrounding areas were cleaned with narrow cotton swabs such that any debris was cleared. After this step, the mice then underwent different procedures depending on whether they were used for awake experiment or for experiments under anesthesia.

#### Electrophysiology in awake behaving animals

We seated a custom spinal clamp implant (Fig. S1A) on the cervical vertebrae and secured it with a clamping screw. While the screw maintained tension, the clamp’s guide posts were cemented in place (Metabond, Parkell) to preserve it. The V-shaped grooves of the implant were cemented to the facet joints, leaving the dorsal surface of the cord accessible. The arm attachments wer cemented to the occipital bone to stabilize the spinal cord and largely eliminate the relative motion between the cord and the brain. After the cement cured, the clamping screw was removed and the skin was sutured over the implant, leaving the access window over the cord exposed, and the window was filled with Kwik-Cast (World Precision Instruments) to keep the cord isolated. For recovery, Mice received extended-release buprenorphine (BPX ER; [dose, route]) and were placed on a heating pad. Once awake, they were transferred to a clean cage and given ibuprofen in the drinking water (0.2 mg/mL, 3 days) and wet food. Mice were single-housed and monitored daily, and showed no apparent impairment in the home cage after a week.

Three weeks after surgery, mice were placed on water restriction following established procedures. Mice received 1 mL of water per day, obtained during head-fixed sessions as pseudo-randomly delivered rewards (7 ul water; poisson distribution with a mean of 100-150s) and supplemented in the home cage after the session if they consumed less than 1 mL. Mice were assessed daily for health. Mice were handled for 1 week and habituated to head fixation over ∼10 sessions, increasing from 10 to 40 min on the platform with pseudo-randomly delivered water, before recording.

After habituation, on the day before the first recording session, the Kwik-Cast was removed and a laminectomy was performed within the access window at the C6 vertebrae under isoflurane anesthesia. The dura was gently removed and replaced with a thin layer of Duragel (Cambridge NeuroTech), which was allowed to cure. The site was then covered with sterile Kwik-Cast. For each recording session, mice were head-fixed, the Kwik-Cast was removed, and a silicon probe (Cambridge Neurotech, 64 Channels, H10) was advanced through the Duragel into the cord with a micromanipulator (Sutter MP285) to a depth of ∼750um at 250um lateral to the dorsal vein. Neural recordings are described below. Sessions lasted approximately 40 min. At the end of each session, the probe was retracted and the site was refilled with Kwik-Cast. Each mouse underwent three recording sessions; recording sites remained viable for up to one week, as judged by visual inspection.

#### Electrophysiology under anesthesia

For experiments under anesthesia, the mice were then given a cocktail of urethane (1.5 g/kg, Sigma) and xylazine (1.2 g/kg, Sigma) for long term anesthesia during the recordings. The isoflurane anesthesia was gradually reduced to zero over the next few minutes. The mouse was then moved to the electrophysiology rig, placed on a heating pad and was then head-fixed on the rig by the headplate. Spinal clamps (Kopf) positioned at C4/C5 were then used to stabilize the spinal cord, and lift the vertebral column up such that the rib cage moved freely during respiration. The right forepaw/hand of the mouse was then attached to the top of the pSCARA manipulandum by applying a cyanoacrylate glue to the glabrous skin. A laminectomy was performed to expose the spinal cord. The dura was then gently removed with fine forceps and spring scissors. To prevent the cord from drying out, the cord was continuously irrigated with saline. Once the dura was removed, silicon electrodes were driven to the appropriate coordinates in the spinal cord and neural responses to movement stimuli were recorded. Recording sessions lasted between 6 to 8 hours. At the end of the recording session, the animal was either perfused and collected for histological analyses or euthanized by cervical dislocation.

For optical tagging experiments in PV^Cre^;Ai32, Calb1^Cre^;Ai32, and Ptf1a^Cre^;Ai32 a laser diode light source (LDFLS_450, Doric Life Sciences) attached to an optical patch cord was used to deliver light to an optical cannula on the surface of the spinal cord (200 μm core diameter; NA = 0.66, Thorlabs). Lbx1^Cre^;Ai32 targets excitatory and inhibitory interneurons, broad activation of these neurons cause widespread activity in the spinal cord that makes optotagging intractable. To overcome these limitations we used the 32 channel μLED Optoelectrodes from the University of Michigan to that can provide localized excitation. To test for light-evoked spiking, at least 50 pulses of light (100 ms duration, 1ms offset ramp) were delivered at 3 different power intensities (5, 10, and 25 mW/mm^2^ at fiber tip) at the beginning and at the end of the recording. Neurons were considered optogenetically identified if they had low jitter (<3ms) and high consistency (>90%) responses to 450-nm light stimulation.

Extracellular recordings in both awake and anesthetized experiments were made acutely using 64-channel silicon probes (ASSY-77 H10, Cambridge Neurotech) or the 32-channel MINT optoelectrode. The voltage signals were amplified, filtered and digitized (16 bit) on a headstage (Intan Technologies, RHD2164), recorded on a 512-channel Intan recording controller (sampled at 20 kHz), and stored for offline analysis.

Extracellular voltage traces were first band pass filtered for 250Hz-7.5kHz. The data were then spike-sorted automatically with Kilosort2.5, and curated manually with Phy2. During manual curation, units containing low-amplitude spikes and/or non-physiological or inconsistent waveform shape were discarded and not included in further analyses. Neurons with fewer than 4 trials in any of the conditions tested were excluded for all analyses. Neural location was estimated from the array’s depth read out from the micromanipulator, and the position of the electrode contact where the neuron had the largest voltage magnitude. Neurons with ISI violations of more than 1% or smaller than 100 μV spike amplitude were excluded from the study. Median (IQR) spike amplitude for the recorded neurons were ∼325 μV ([250 410]) and Median (IQR) ISI was 0.1% ([0 0.4%]).

### Kinematic tracking

Hand kinematics during awake behavior were tracked by using synchronized video captured at 250Hz from 4 cameras (Basler, Ace 720-520um) with overlapping view angles. The cameras were calibrated using a checkerboard to establish the intrinsic and extrinsic parameters for the cameras. SLEAP was used to perform key-point tracking on the hand. Separate models were trained for each view, and the performance was improved by selective retraining of videoframes where the reprojection errors were high. The kinematics were then filtered using a low-pass 50Hz 8 pole Butterworth filter.

### Tuning curves and Spatial rate maps

To visualize how firing relates to movement, we computed tuning curves and spatial rate maps from the z-scored firing rate of each neuron. For tuning curves, each kinematic variable (position along x, y and z, and speed) was divided into 20 bins containing equal numbers of samples, so that every bin was well sampled. Bins with fewer than 20 samples were discarded. Error shading on each bin’s mean firing rate were computed using a moving-block bootstrap: the rate time series was resampled in contiguous blocks rather than as independent samples, to preserve local temporal autocorrelation structure; each bin’s mean was recomputed across 200 such resamples, and the standard deviation of the resulting bootstrap distribution was taken as the bin’s standard error.

Spatial firing-rate maps were computed by projecting 3D hand/limb position onto the plane defined by its first two principal components, chosen to minimize fragmentation of the trajectory relative to a fixed cardinal plane. Position within this plane was binned into a 2D histogram (30 × 30 equal-width bins), and for each bin, occupancy (sample count) and mean firing rate were computed; bins with occupancy below a minimum threshold (5 samples) were excluded (mean rate set to NaN) to avoid unreliable rate estimates from sparsely sampled locations.

### Neural Encoders

To determine the amount of kinematic information carried by cervical spinal cord neurons during awake spontaneous behavior, we used four major classes of the neural decoders: linear models, non-linear additive models, gradient-boosting trees and gated recurrent units.

For the models, we used the forelimb position tracked using video (described in Kinematic tracking) and spikes were counted in 20 ms bins. Spike counts were smoothed with a Gaussian kernel (σ = 40 ms) and z-scored for each neuron, except for the Poisson GLM, which was fit to raw counts. All models used the same 11 features computed from the tracked position: position (x, y, z), velocity (vx, vy, vz), speed, acceleration (ax, ay, az) and acceleration magnitude.

To account for the strong autocorrelation of movement and firing, we used block cross-validation. Each recording was divided into 64 contiguous blocks, which were assigned in alternation to three folds so that every fold sampled the whole session. Models were trained on two folds and scored on the third by the held-out coefficient of determination (R^2^), averaged across folds.

We compared four model classes of increasing flexibility. The linear model was a Poisson generalized linear model (GLM) with a log link and L2 regularization. The additive model was a generalized additive model (GAM) in which each feature contributes a smooth, independently fit function (a cubic spline), with no interactions. We used it to estimate tuning curves and each feature’s relative contribution, defined as the variance of its fitted function as a percentage of the total across features. The interaction model was gradient-boosted regression trees, which capture nonlinear effects and interactions between features. The recurrent model was a gated recurrent unit (GRU) network of one layer with 32 hidden units that received the preceding 500 ms of the same features, to test whether movement history adds information beyond the instantaneous state.

Analyses were performed in MATLAB 2021b.

### pSCARA Motion Control System and Movement Stimuli

We built a robotic manipulandum based on the parallel selective compliance articulated robot assembly (pSCARA) designed for rodents. The robot has two degrees of freedom, and was driven by two direct current motors (Maxon RE-max 21) and coupled with two high-resolution optical rotary encoders (Gurley Precision Instruments R120B). The motors were driven with dedicated DC driver boards (VNH5019 Pololu Inc.) and the current drawn was measured with hall-effect based (ACS724) high resolution current sensors (800mV/A).

The control system was a single board real time i/o field programmable gate array (sbRIO-9637, NI). The sbRIO is an integrated system that contains a real-time controller and an FPGA. Similar to previously described systems we used a nested control system. The lowest level of the control a standard proportional-integrative-derivative (PID) feedback controller operating at 1 kHz was implemented on the FPGA to control the motors. The second level of control was implemented on the real time controller which performed kinematic transformations from the coordinates of the robot joints to Cartesian coordinates. The final level of control was on the interfaced personal computer running LabView which determined the stimulus type and was responsible for writing the acquired data to disk. We sampled the manipulandum’s kinematics at 1 kHz.

We fine-tuned the PID values and validated the motion control system by the comparing the commanded kinematics to that executed by the motors when it was attached to a mouse forelimb (Fig. S1A-C). This robotic system allows for delivery of stimuli with arbitrary kinematics. We chose stimuli that mimicked center-out reaching movements with minimum-jerk dynamics. The range of speeds (20-80 mm/s) was chosen to be consistent with those observed in previous behavioral studies. Executed trajectories were read out from optical rotary encoders on the motors and compared with the commanded trajectories over time for all 64 directions at three speeds, with 10 trials each (1,920 movements; Fig. S1B).

### Electrophysiological Analysis During Anesthesia

For this study we recorded a total 3273 neurons from 41 mice. For each neuron, a peristimulus time histogram (PSTH) was constructed for each direction of the movement stimulus. The PSTH was constructed by including spikes from 50ms before the onset of the movement stimulus to 200 ms after the onset of movement stimulus, with time bins of 5ms. The PSTH was then filtered with a 25ms Gaussian window.

To perform unsupervised clustering of the neural responses, we first constructed the feature vector by horizontally concatenating the PSTH for each stimulus direction in clockwise order, starting with the direction of the peak firing rate. This feature vector was then z-scored and then vertically concatenated to create a 2D matrix of size N x (D x T). Where N is the number of neurons, D is the number of stimulus directions and the T is length of the PSTH vector. For the analyses in Figure 1, this was 1570 x (64 x 50). UMAP dimensional reduction was then applied to this matrix to get the representation in Figure 1. The UMAP transformation revealed three distinct clusters that could easily be identified with a simple watershed algorithm.

For modelling the single neuron firing rates for the Gain-field neurons we tested two gain field models both of which took a similar form:

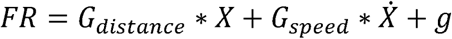

Where, *FR* is the firing rate calculated as the PSTH, *X* is distance and *X* is instantaneous speed of the hand.

The gain-field models differ in dependence of the gain-function on angle. The cosine gain-functions have the form:

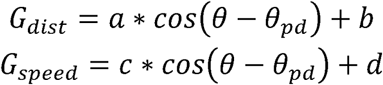

The von Mises gain function has the form:

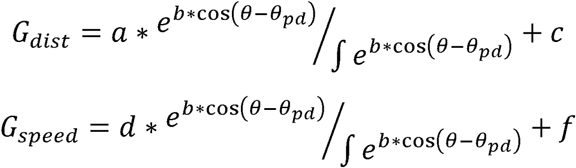

*θ* is the movement angle and *θ_pd_* is the preferred direction. *a,b,c,d,f* and *θ_pd_* are parameters to be fit. We fit the models using the ‘fminsearch’ function in MATLAB 2021b.

Speed/Position ratios were calculated at the ratio of G_speed_ and G_dist_ estimated from the fit.

Firing fields for neurons were calculated by generating a 2D spatial histogram quantifying the number of spikes that occurred in each spatial bin over all the trials. The 2D histogram had a bin size of 200μm x 200μm. The firing fields were then normalized to the 99^th^ percentile of number of spikes in a bin.

To estimate the tiling of space, firing field maps were z-scored and were binarized. Significant bins (zscore > 1.65) were given the value 1 and others were assigned 0. This binarized firing field of each cell was then overlaid and normalized by the total number of neurons to calculate the tiling shown in Fig. 1J.

For the dPCA analyses the design matrix was constructed as N x D x S x T. Where N is the number of neurons, D is the number of stimulus directions, S is the different speeds and the T is length of the PSTH vector. For the analyses in Figure 2, this was 138 x (64 x 3 x 50). The dPCA decomposition was performed using the code made available by the original study (Kobak et al 2016). For Fig. S6B-C, the neural data for the independent conditions are projected into the dPCA space estimated on the full dataset. For Fig. S6D-F, the dPCA was calculated independently for each of the ‘Clusters’ and the combined neural activity was plotted in the top 3 condition-independent dimensions identified for each dPCA calculation.

### Neural Decoders

To estimate limb kinematics from neural activity we used three classes of models: Linear models, maximum likelihood models and recurrent neural networks.

The linear models took the form of

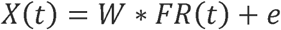

Where, X(t) is the x,y position of the hand kinematics at time t, W is the learned weights, FR is either the binned or the binned and smoothed firing rate of the neurons, and e is the estimation error. For cross-validation of the models, W was calculated by using half the trials to calculate the FR and then performing a pseudoinverse. The model was then tested on the mean FR estimated from the held out trials, the mean absolute error and the r-squared for the cross validation was calculated on these predictions. For single trial predictions, all the trials except one were used to calculate the mean FR and estimate W. The model was then tested on the remaining trial, and the mean absolute error and r-squared were calculated from this test.

The linear models performed poorly on binned neural data (Fig. S7A, top), although their performance improved with smoothed firing rates (Fig. S7B, top). These models were sensitive to the number of neurons used to make the kinematic predictions (Fig. S7A,B) and showed improvement with larger neural populations. However, the single trial decoding errors with the full population of neurons were still substantial (∼800 um) and provided poor estimates of the moment-by-moment kinematics (Fig. 2C-E and Fig. S7A,B bottom).

The maximum likelihood models (MLE) used probabilistic priors of the ensemble neural activity to estimate the kinematics in the current moment. To validate the models we used half the trials to generate kinematics priors for neural firing patterns (FR_train_), and then other half to generate spike probabilities (FR_test_). As in the linear models, we evaluated these probabilities for both binned and binned+smoothed data. For decoding, we find the closest match of the FR_test_ vector to the FR_train_ vector in each time step. The kinematics of the most similar FR_train_ vector is then determined as the decoded kinematic state, this can be formalized as:

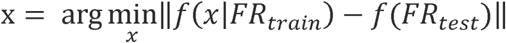

Similar to linear models, the maximum likelihood models showed improved prediction of the kinematics with smoothed firing rates and larger neural populations (Fig. S7C,D), and had substantially improved performance compared to the linear models (Fig. 2C-E and Fig. S7C,D bottom).

For the recurrent neural network models, we used the LSTM architecture. LSTM cells were implemented as described in the original literature. We only needed one LSTM layer to estimate kinematics with high accuracy. Network size was varied by changing the number of hidden units in the layer. Networks were trained till the validation loss plateaued (typically 200 epochs). To generate training data for the LSTM networks, we created a single trial pseudopopulation data set. We did this by first randomly selecting a subset of neurons, and for then for each neuron randomly selecting one trial for each direction. One trial was held out for each neuron and each condition, and wasn’t used in the training dataset. We repeated this ∼1000 times to get a training dataset. This led to a dataset of N x D x FR x 1000, where N is the number of neurons used for the training set, D is the number of stimulus directions (typically 64, but varied for experiment in Fig. S7G) and FR is the spike count for that neuron in that trial, binned with 5ms. The test set was created similarly by using the held out data, the test set always had the dimensions N x D x FR x 1.

Since LSTM networks have a large set of parameters that can lead to ‘memorizing’ the data, we undertook a series of validations to confirm that the network was learning a transformation. First, we tested the kinematic predictions on stimulus directions that were held out of the training data. If the LSTM was trained on at least ¼ of the stimuli, its predictions were entirely unimpaired (Fig. S7G). Additionally, an LSTM should perform very well on shuffled data if it is simply memorizing the data, but very poorly if it is learning a transformation. We found that LSTM networks performed poorly on shuffled data (Fig. S8G).

The bin width for all models was 5ms and the firing rate vectors were smoothed with a 25ms wide Gaussian window.

### Tamoxifen induction

Adult *Avil^CreERT2^;MafA^FlpO^;Tau^ds-DTR^; R26^ds-tdT^* mice (>8 weeks old) were administered tamoxifen (100 mg/kg, intraperitoneally) once daily for 5 consecutive days to induce CreERT2-mediated recombination. Tamoxifen was prepared in corn oil at a final concentration of 20 mg/ml. Experiments were performed at least 3 weeks after the final injection to allow enough time for Cre-mediated recombination and subsequent expression of recombined alleles.

### Immunohistochemistry

Mice were anesthetized by a single intraperitoneal (i.p.) injection (10 ml/g body weight) of solution containing 10 mg/ml ketamine and 1 mg/ml xylazine immediately prior to perfusion with 20 mL of 4% paraformaldehyde in PBS. Spinal cords and DRGs were dissected and post fixed for 1 hr at room temperature (RT), then rinsed 3 times in PBS and cryoprotected in 30% sucrose-PBS overnight at 4C. Spinal cords were embedded in OCT (Tissue-Tek) and cryosectioned at 40μm using a Leica CM3050 cryostat, DRGs were similarly cyrosectioned at 14μm. Sections were dried at RT and stored at −20C. Before staining OCT was removed with a 15 min PBS wash. Slides with sections were then incubated in Blocking Solution (PBS, 10% donkey serum and 0.3% Triton X-100) for 1-2 hr at RT and then incubated with the primary antibody diluted in the antibody solution (PBS, 1% donkey serum and 0.3% Triton X-100) overnight at 4C. Sections were then washed 3 times (15 min each) in PBT (PBS and 0.1% Triton X-100) before being incubated for 2 hr at RT with antibody solution containing donkey-raised fluorophore-conjugated secondary antibodies (1:1000; Jackson Laboratories). Sections were again washed 3 times (15 min each) in PBT before being mounted with Aqua-Poly/Mount (Polysciences). A Zeiss LSM700 or an LSM900 confocal microscope was used to capture images. ImageJ software was used to count neurons, the Cell Count plug-in was used to analyze co-localization.

### Neuronal Ablation

Neuronal ablations for the experiments related to Fig. 5 were performed by two i.p. injections of diphtheria toxin (DT, 50 ng/gram of weight; List Biological Laboratories) three days apart in animals >16 weeks of age. Electrophysiological experiments for recording movement responsive neurons were then performed soon after (<3 weeks). For the behavioral experiments the animals received a single i.p. injection with the same total dose (100 ng/gram of weight)

### Drug Administration

Clozapine-N-oxide (Sigma) was dissolved in DMSO and then diluted with 0.9% sterile saline such that the concentration of DMSO did not exceed 1% of the injected solutions. For experiments related to the electrophysiological validation of the chemogenetic silencing (R26^ds-tdT-hM4D^) mice were give i.p. injections of saline before movement stimuli were delivered. After the saline responses were acquired, animals were injected with CNO (3 mg/kg). Approximately 10-20 minutes after the CNO injection, the second set of movement stimuli were delivered. For the behavioral experiments in Fig. 4, animals were given saline on day 1, CNO on day 2 and saline again on day 3.

### Joystick Center-out Reach Task

We trained mice to use a joystick to make center out reaches to specified targets in an automated home cage training system. Individually housed mice were trained to enter a nose port, grab a fixed post with their left forelimb, and then grasp the joystick with their right forelimb. The mice were then trained to make rapid reaching movements to a spatial target 4 mm out from the center. All mice learn to reach to the target within 10 days of training. After the animals were consistently reaching out to a target, we started restricting the amount of time mice spend in home cage to 3 hours. Since the only source of water for these mice is through this task, the animals had to perform enough successful trials to meet their water needs within this time frame. After 3 days of training with the time restriction, we collect three more days of baseline trajectories. For the neuronal ablation experiments, on the fourth day mice received an injection of DT. The mice were then tested with the same time restriction for three more days. For the hM4D experiments, mice were given an i.p. injection and placed into the home cage 10 minutes after the injection. Mice given saline on the fourth day, CNO on the fifth day and saline again on the sixth day. The time restriction was necessary to make appropriate comparisons between saline and CNO trials since CNO’s potency is time dependent with maximal effects limited to within a few hours of the infusion.

### Analysis of Joystick Trajectories

Trajectories were acquired at 1 kHz and low-pass filtered at 50 Hz with an 8-pole Butterworth filter in software. The reach direction was defined as the angle at which the target distance was transected by the trajectory. Velocity was calculated as a one sample difference of the filtered position vector. To make sure that we were specifically analyzing trajectories that were attributable to right forelimb movement with the animal in a consistent posture, only the trajectories that were contacted after the nose poke and fixed-post contact were considered valid trials eligible for reward. If the mouse exited the nose poke, lost contact with the fixed post or the joystick, the trial was immediately failed. The mouse also failed the trial if it reached in the wrong direction. There were no cues for failed or successful rewards (except the water delivery apparatus). However, a blue LED masking light roughly at eye level to the mouse was turned on at joystick contact. The mouse had to re-contact the joystick to start a new trial after both successful and failed trials.

A reaching movement typically consists of initial reach segments followed by corrective segments. To identify the initial reach end-point points of the reaching movements we decomposed complex trajectories into submovements, where submovement boundaries were defined by temporally coincident minima of velocity and radius of curvature. The end-point of the first reach segment to cross the 1 mm distance from the center was defined as the initial reach end-point.

### Statistical Analysis

All statistical analyses were performed in Matlab 2021b. Neurons were considered modulated by the movement stimulus if the number of spikes after movement onset were significantly different than baseline in any of the stimulus directions. Significance was assessed by performing a t-test for the mean number of spikes before and after the stimulus, with a p<0.01. Statistical analyses for measures of kinematics were performed using a hierarchical bootstrap for mice, sessions and trials. We used a two-tailed t-test for assessing difference in mean cell counts of histological analyses.

R^2^ values are reported for the neural encoding models (Fig. 1), the gain-field models (Fig. 2), dPCA axes (Fig. 4) and for neural decoding (Fig. S8). An R^2^ value was reported only if its bootstrap-derived p-value was below 1 × 10□□.

## Acknowledgments

We thank Akira Nagamori for help with 3D tracking of the hand position in the awake animals. We thank Jose Roberto Lopez Ruiz and Euisik Yoon at the Univeristy of Michigan for access to- and training for the μLED probes supported by NSF NeuroNex MINT Hub (Award 1707316). We thank Graziana Gatto for advice on selection of mouse models in the early planning stages of this work. We thank Nehal Bokka for technical assistance with histology. We thank Eiman Azim, Ruidong Chen, Joe Fetcho, Jesse Goldberg, Kee Wui Huang, Mayank Mehta and Akira Nagamori for feedback and comments on the manuscript.

## Author Contributions

Conceptualization: TB, MDG

Methodology: TB

Investigation: TB

Writing – original draft: TB

Writing – review and editing: TB, MDG

## Data and materials availability

All data, code, and materials used in the analysis will be made available by TB on reasonable request.

## Supplementary Materials

Materials and Methods

Figs. S1 to S11

Movie S1

Movie S2

**Fig. S1.**
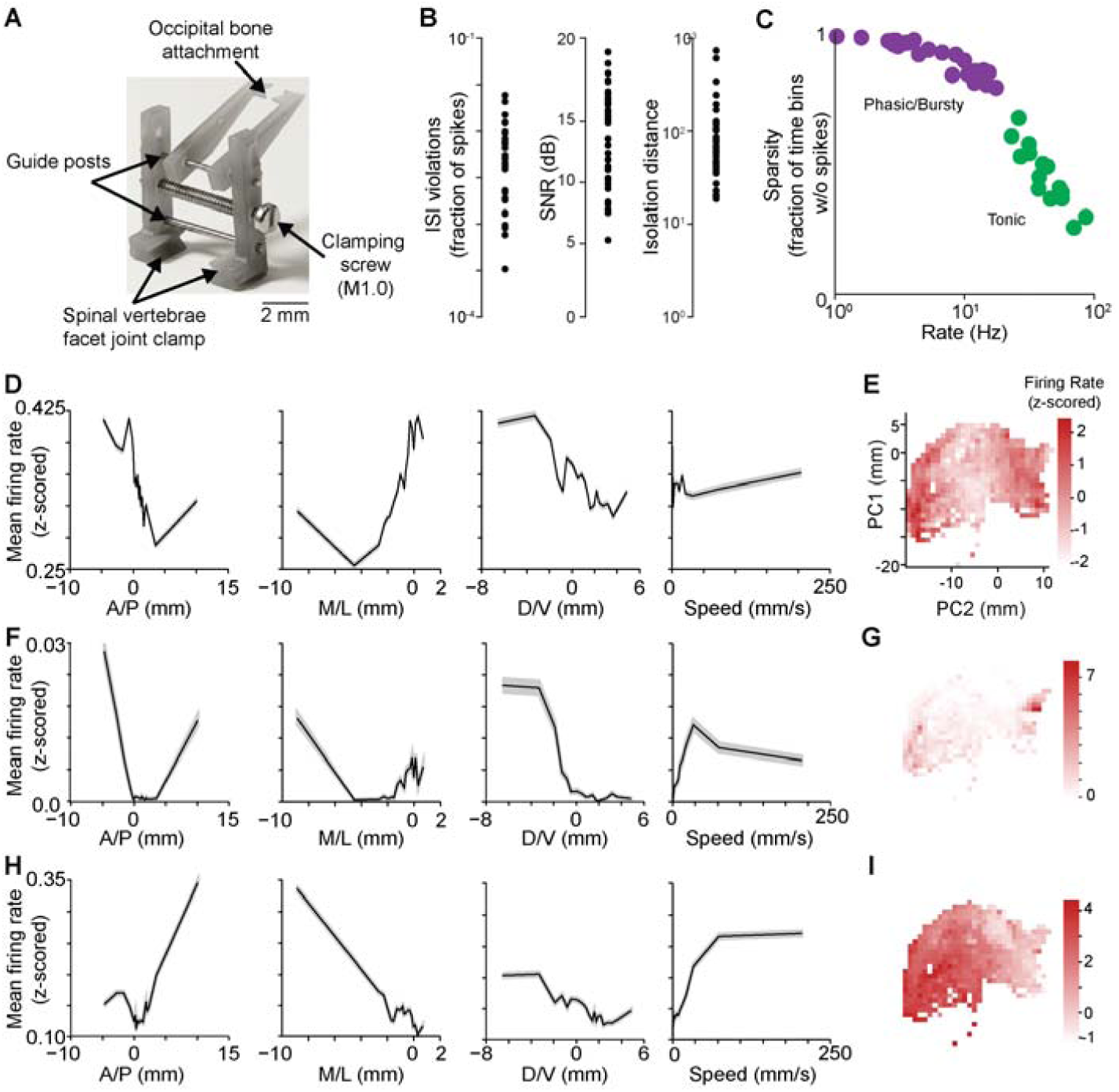
Cervical spinal cord neurons recorded in awake, behaving animals were well isolated and showed diverse discharge patterns, kinematic tuning and spatial firing. **(A)** An implant to stabilize the cervical spinal cord for acute electrophysiology in awake behaving animals by reducing motion relative to the skull. **(B)** Neurons recorded in the cervical spinal cord in awake behaving animals were well isolated with good signal-to-noise ratio. **(C)** Neurons had varying response properties **(D)** Firing rate of an example neuron as a function of the hand position in anterior-posterior, medial-lateral and dorsal-ventral axis, and as a function of hand speed. **(E)** Spatial firing-rate map for example neuron from (D). Hand position was projected on a plane defined by the first two principal components of the 3D position, chosen to minimize the fragmentation of the trajectory relative to the A/P, M/L and D/V cardinal planes. **(F-I)** Same as (D-E) for two other example neurons. Rates are z-scored relative to the neuron’s own session mean

**Fig. S2.**
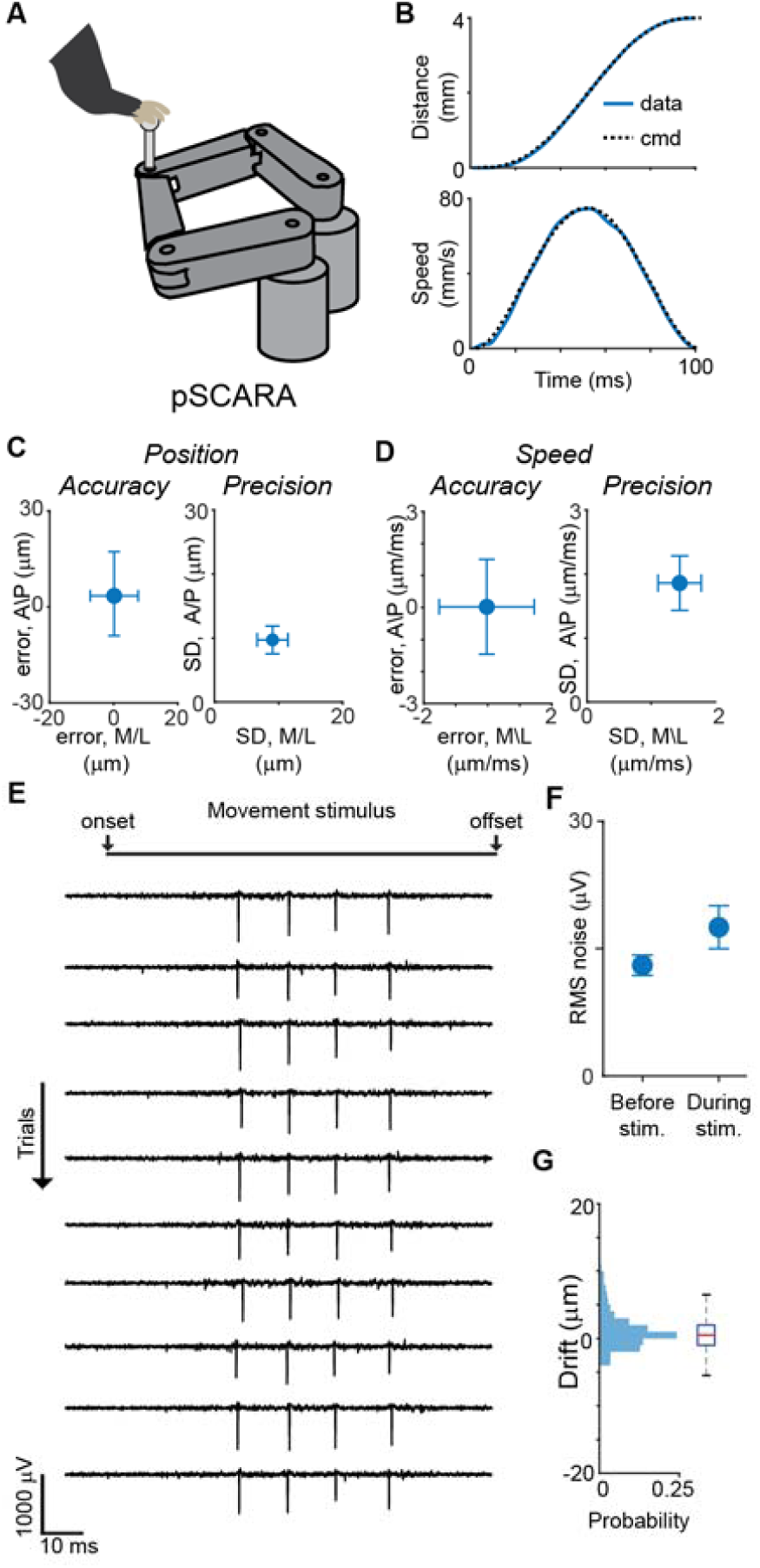
Stable in vivo electrophysiological recordings from the cervical spinal cord while delivering precise motion stimuli to the forelimb. **(A)** Schematic of a mouse forelimb attached to a parallel selective compliance assembly robot arm. **(B)** Example temporal profiles of the robot’s speed and position. The comparison of commanded (dashed black line) between and observed (blue line) kinematics illustrates the precision of the robot. **(C)** Quantification of the accuracy and precision of the robot’s position relative to the commanded position. Values are represented as Mean and standard deviation. **(D)** Same as (C), for speed. **(E)** Example filtered and whitened electrophysiological traces for a single channel as a movement stimulus was delivered to the mouse forelimb. Ten trials were recorded ∼ 5 mins apart from each other. Note the minimal effect of movement stimuli on the electrophysiology and the temporal stability of the neural spike amplitudes. **(F)** Quantification of the noise induced by the delivery of movement stimulus to the forelimb, represented as mean +/- standard deviation. **(G)** Estimated drift of the neural spikes over a ∼6 hour recording session. Red line is the median, solid blue lines are the 25-75^th^ percentiles, and whiskers are 1.5 times the interquartile range.

**Fig. S3.**
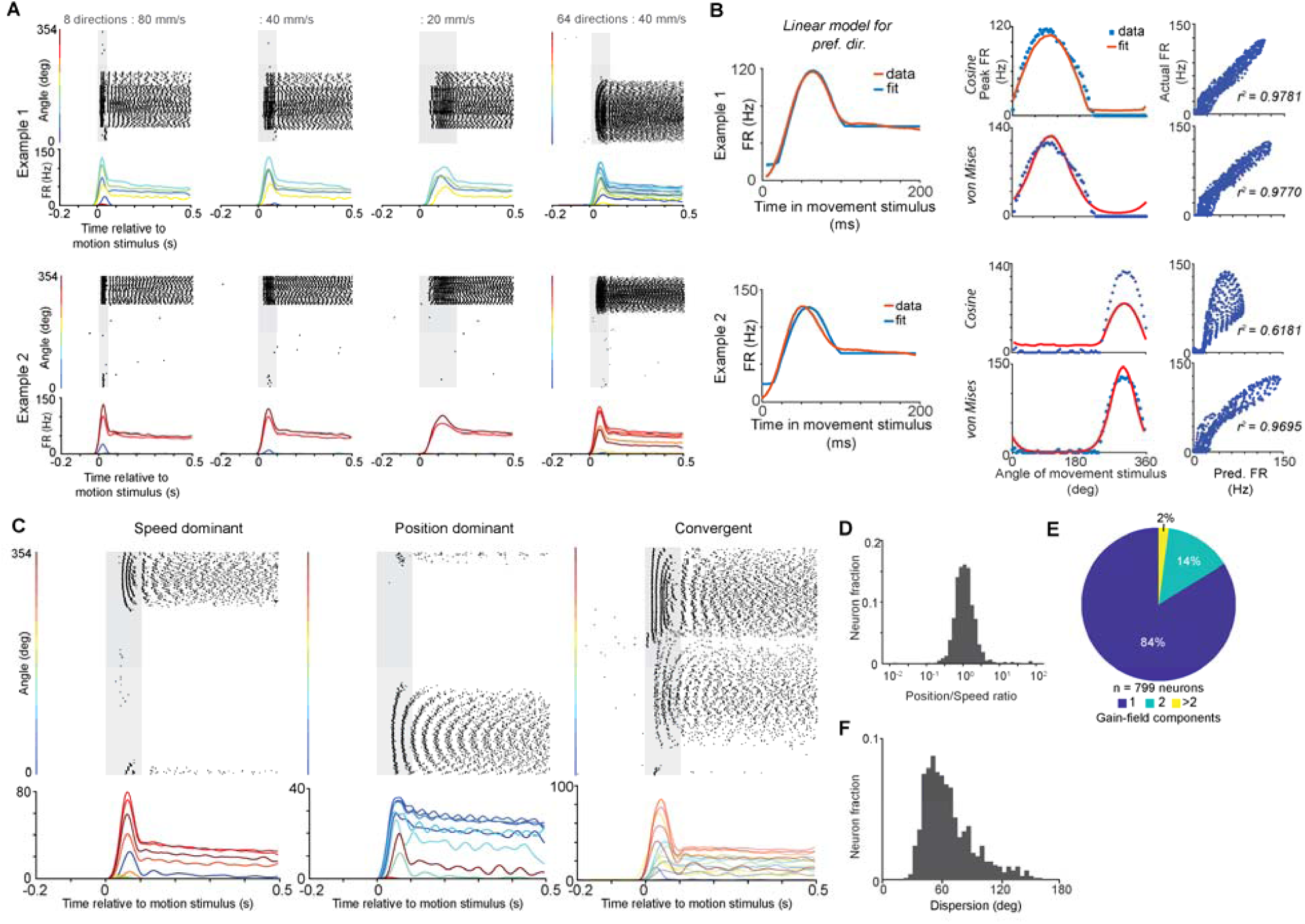
Firing rates of ‘Cluster 1’ neurons are modulated by speed, position and direction of movement stimuli. **(A).** Rasters and rate histograms of two example neurons across different speeds and directions. **(B).** Fits of the firing rates of the example neurons in (A) to a linear gain-field model where firing rates are dependent on speed, direction and position. Compared to cosine tuning, the flexibility in the tuning width of von Mises models enables better fits in the ‘Gain-field’ neural population. **(C).** Gain-field neurons can be differentially sensitive to speed (left) and position (right). A few Gain-field neurons were tuned to multiple movement directions (right). **(D).** The relative sensitivity to position and speed of Gain-field neurons is normally distributed. **(E).** The vast majority of Gain-field neurons have single peaked directional tuning. **(F).** Tuning widths for individual Gain-field neurons.

**Fig. S4.**
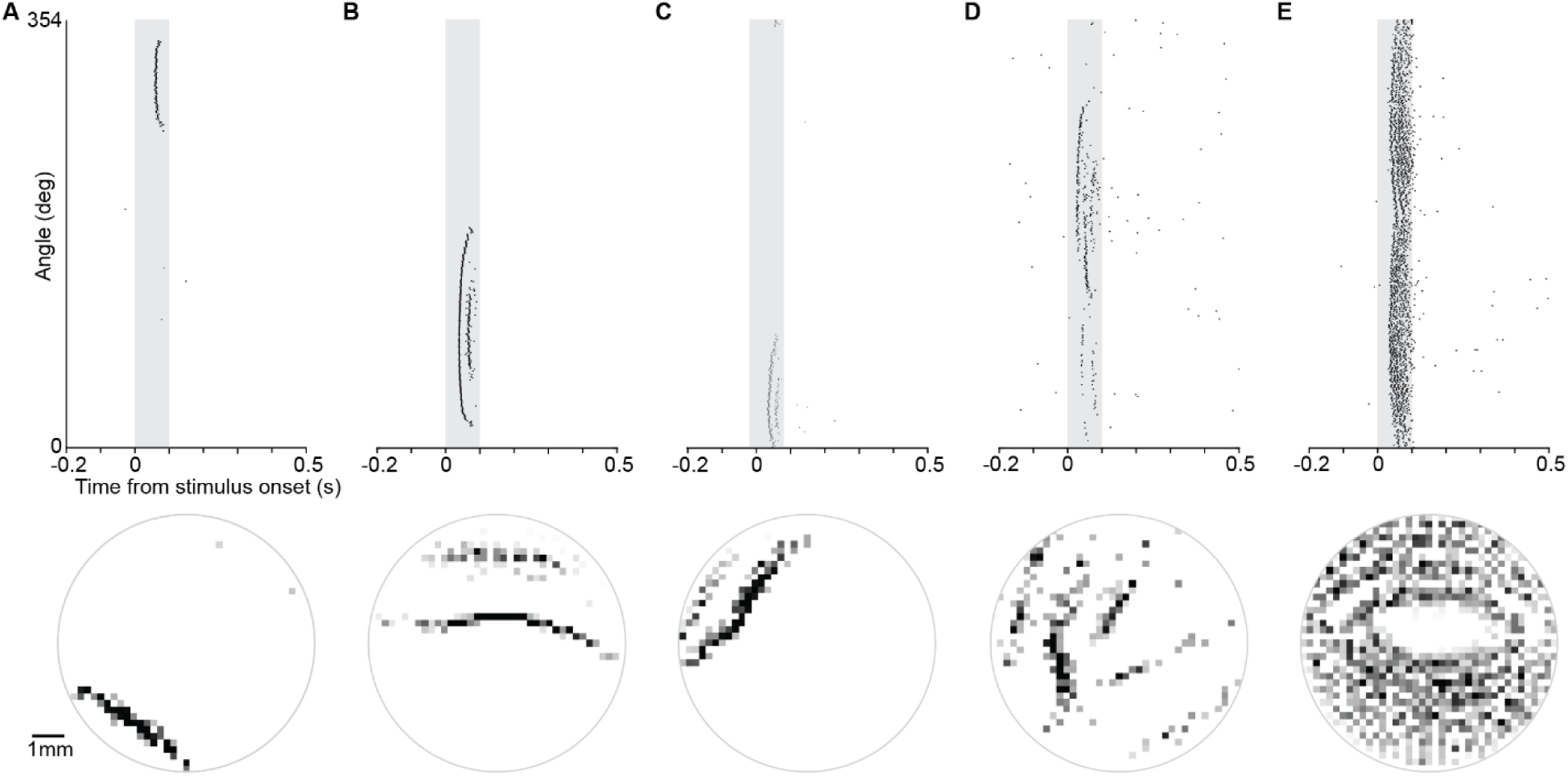
Firing patterns of sparse-coding neurons have multi-scale coverage of the kinematic state space. **(A-E)**. Rasters (top) and spatial firing fields (bottom) of five example neurons from Cluster 2, i.e. Phasic neurons. The examples illustrate the range of spatial firing specificity observed in this cluster.

**Fig. S5.**
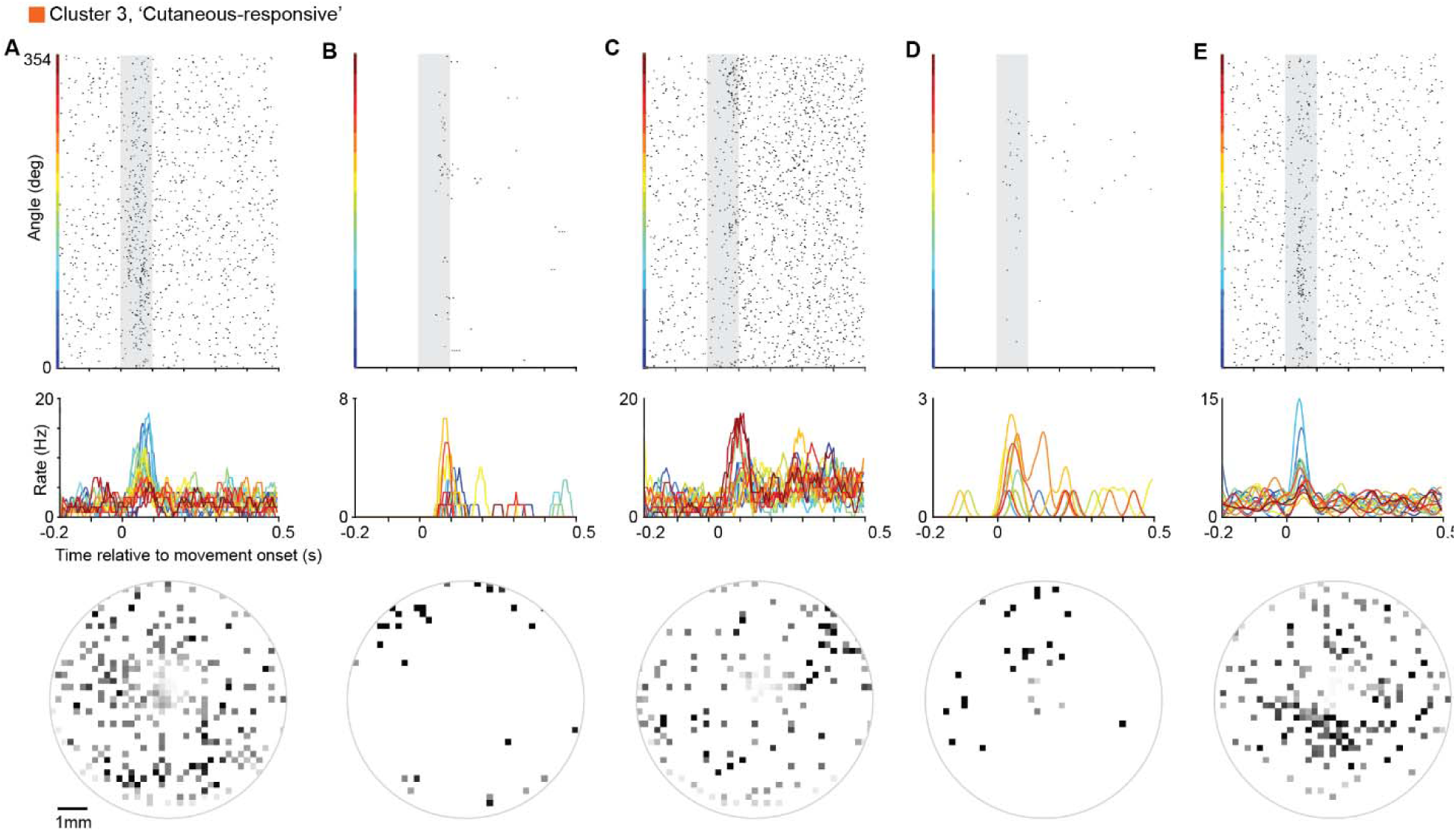
Firing rate modulation of Cluster 3 neurons in response to movement stimuli is inconsistent and spatially unstructured. **(A-E)**. Rasters (top), rate histograms (middle) and spatial firing fields (bottom) of five example neurons from Cluster 3, the Cutaneous-responsive cluster. The examples illustrate that although the neural firing rates of these neurons is significantly modulated by movement stimuli, the responses are inconsistent across trials and unstructured in space

**Fig. S6.**
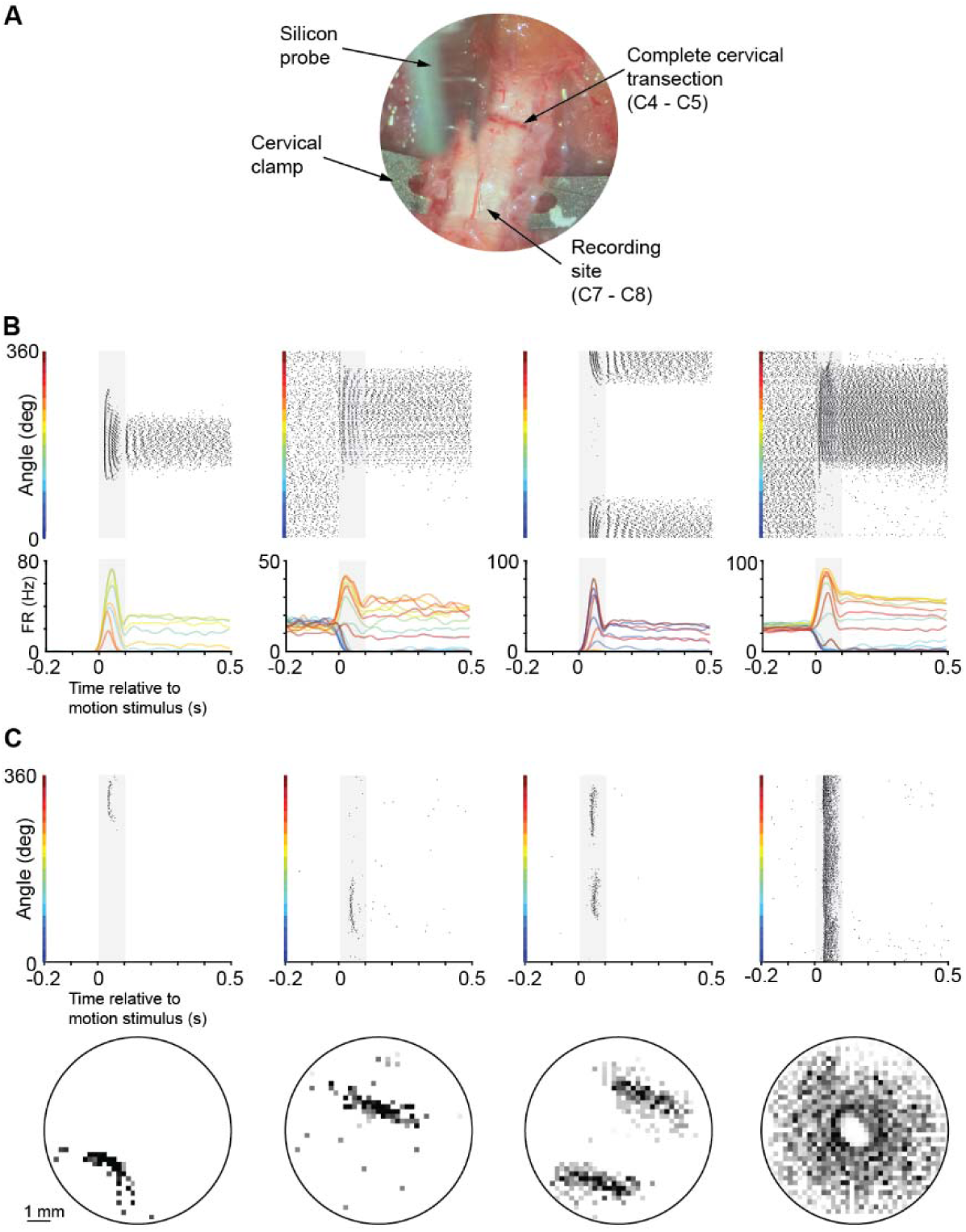
Neural code for forelimb proprioception in the spinal cord persists after a complete transection between the spinal cord and the brain. **(A).** Image of electrophysiological recordings from the C7-C8 segment of cervical spinal cord after a complete transection at the C4-C5 segment. **(B).** Rasters and rate histograms of four example ‘Gain-field’ neurons recorded from the cervical spinal cord after transection. **(C).** Rasters and spatial firing fields of four example Phasic neurons recorded from the cervical spinal cord after transection.

**Fig. S7.**
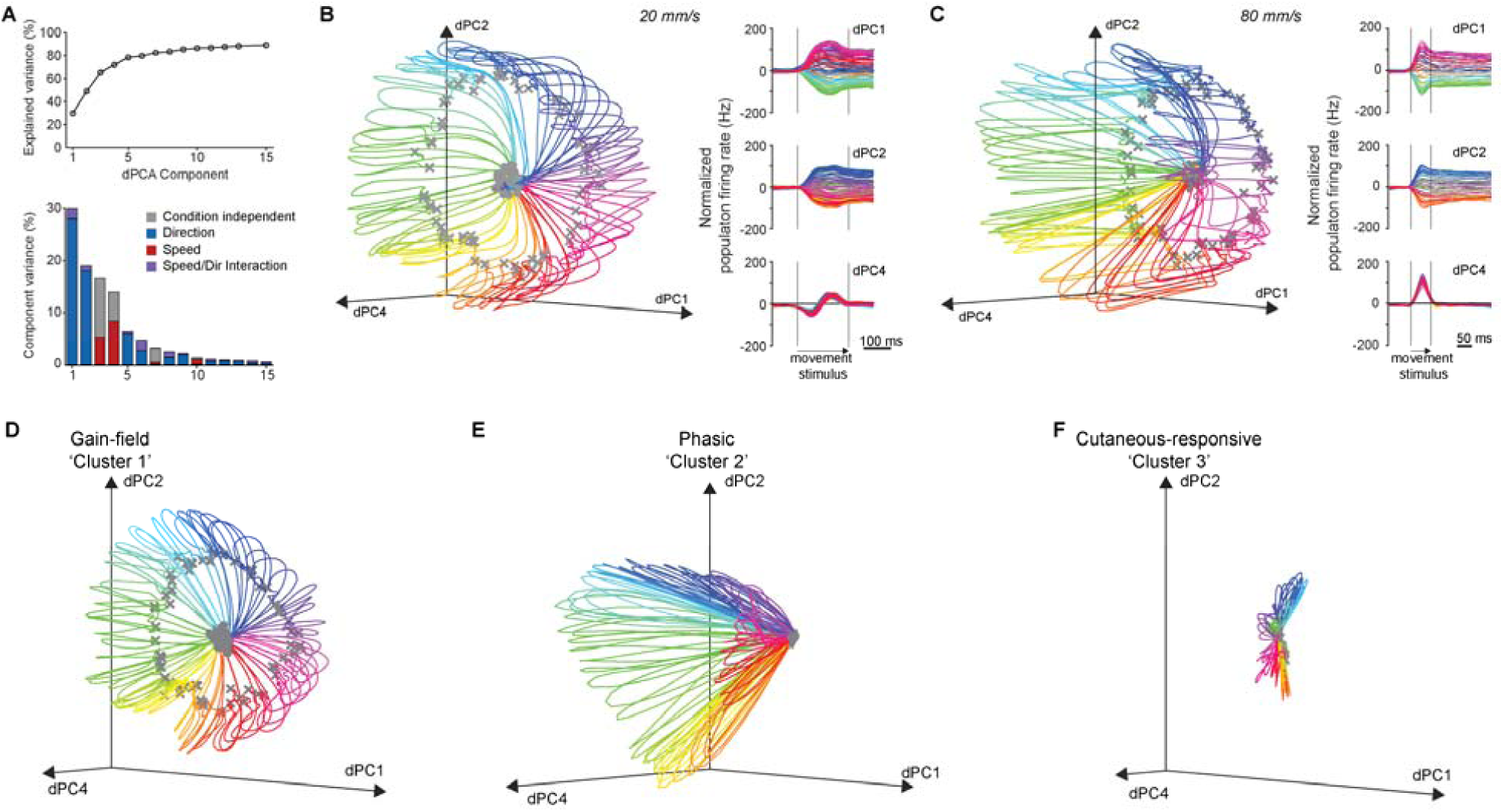
Neural geometry for proprioception in the cervical spinal cord is consistent across different peak speeds, and is constructed by the Gain-field and Phasic neurons. **(A).** The first five demixed principal components (dPCs) can describe ∼80% of the variance in neural activity in the cervical spinal cord evoked by a movement stimulus (top). dPCs were labelled by the relative contributions of the direction and speed conditions to the variance (bottom). **(B-C).** The structure of the neural geometry is invariant to speed. The dynamics of the neural population when the average limb speed was 20 mm/s (B) and 80 mm/s (C) maintains the ordered representation of speed and direction. **(D-F).** Neural clusters contribute differentially to the construction of the neural geometry. The Gain-field (D) and Phasic neurons (E) contribute to the position and speed components, while the Cutaneous-responsive neurons (F) do not form an ordered structure.

**Fig. S8.**
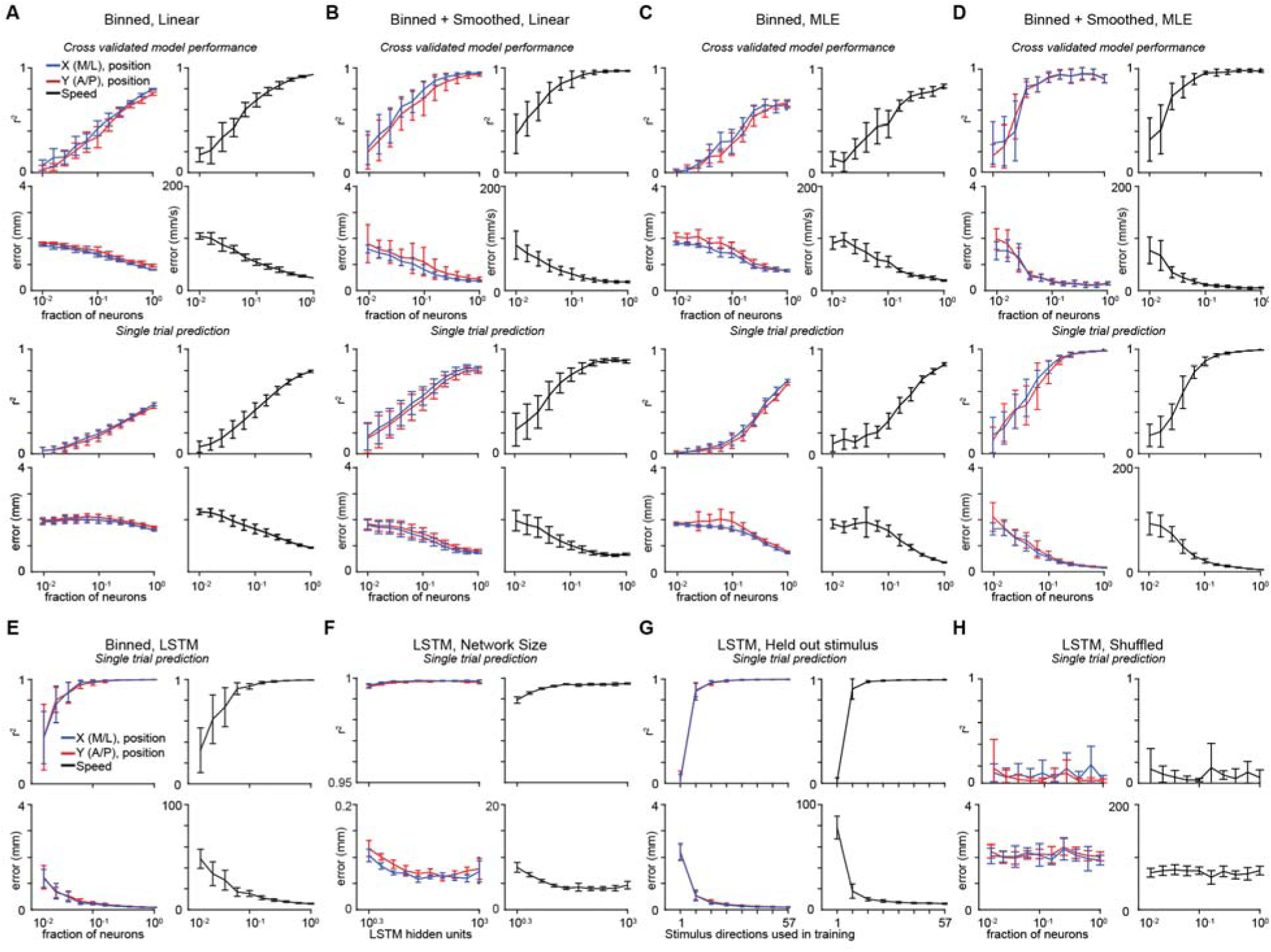
Linear, maximum-likelihood and recurrent neural network models can estimate moment-by-moment kinematics of the forelimb from neural responses in the cervical spinal cord. **(A-B).** Cross validation (top) and single trial predictions (bottom) of hand kinematics from binned (A) and smoothed (B) neural data when using a linear model on different fractions of the neural population. **(C-D).** Plotted as in (A-B), but for maximum-likelihood estimate (MLE) models. **(E).** Plotted as in (A) but for long-short term memory recurrent neural network models. **(F).** Increasing LSTM network size beyond 8 hidden units has a minimal effect in increasing model accuracy. **(G).** LSTM networks can robustly estimate hand kinematics in trials with stimuli they were not trained on. **(H).** LSTM network performs poorly when the identity of the trials is shuffled, demonstrating that this network is not overfitting and memorizing neural patterns for individual trials. Values are plotted as mean +/- standard deviation.

**Fig. S9.**
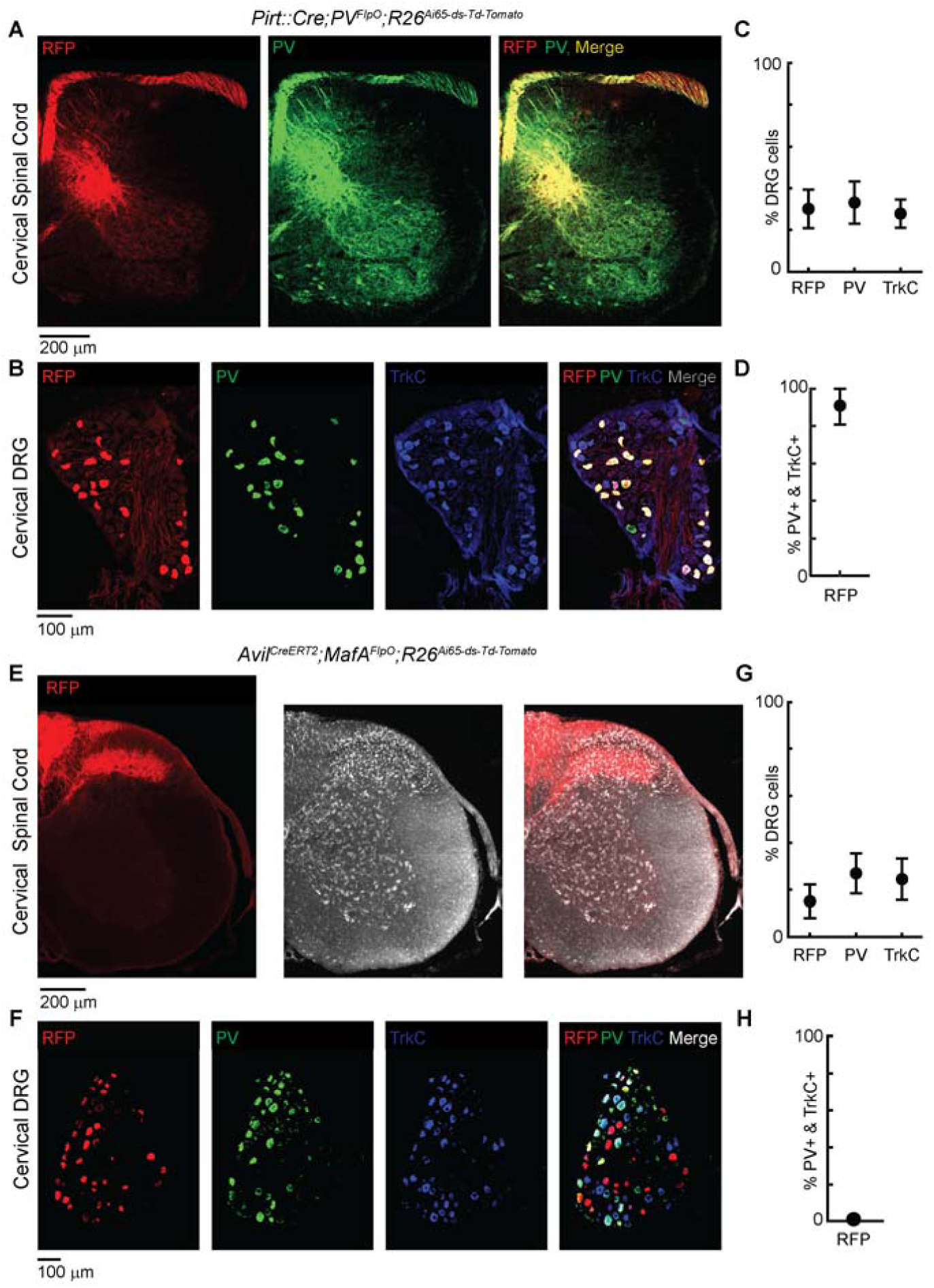
An intersectional strategy to selectively target muscle, tendon and cutaneous afferents. **(A-D).** Immunohistological validation of the intersectional genetics strategy to specifically target muscle afferents. Representative images of the cervical spinal cord (A) and a cervical dorsal root ganglion (B) from a *Pirt::Cre; PV^Flpo^; R26^ds-tdT^*animal (10 days postnatal). This intersection doesn’t target any neurons in the central nervous system, and captures ∼90% of the muscle afferents (D) which are identified by co-expression of TrkC and PV. **(E-H).** Immunohistological validation of the intersectional genetics strategy to specifically target cutaneous LTMRs. *Avil^CreERT2^;MafA^FlpO^; R26^ds-tdT^* animal (>56 days postnatal). This intersection shows no overlap with muscle afferents in the spinal projections patterns (E) and has no overlap with the muscle afferents in the DRG (H). Values are plotted as mean +/- standard deviation.

**Fig. S10.**
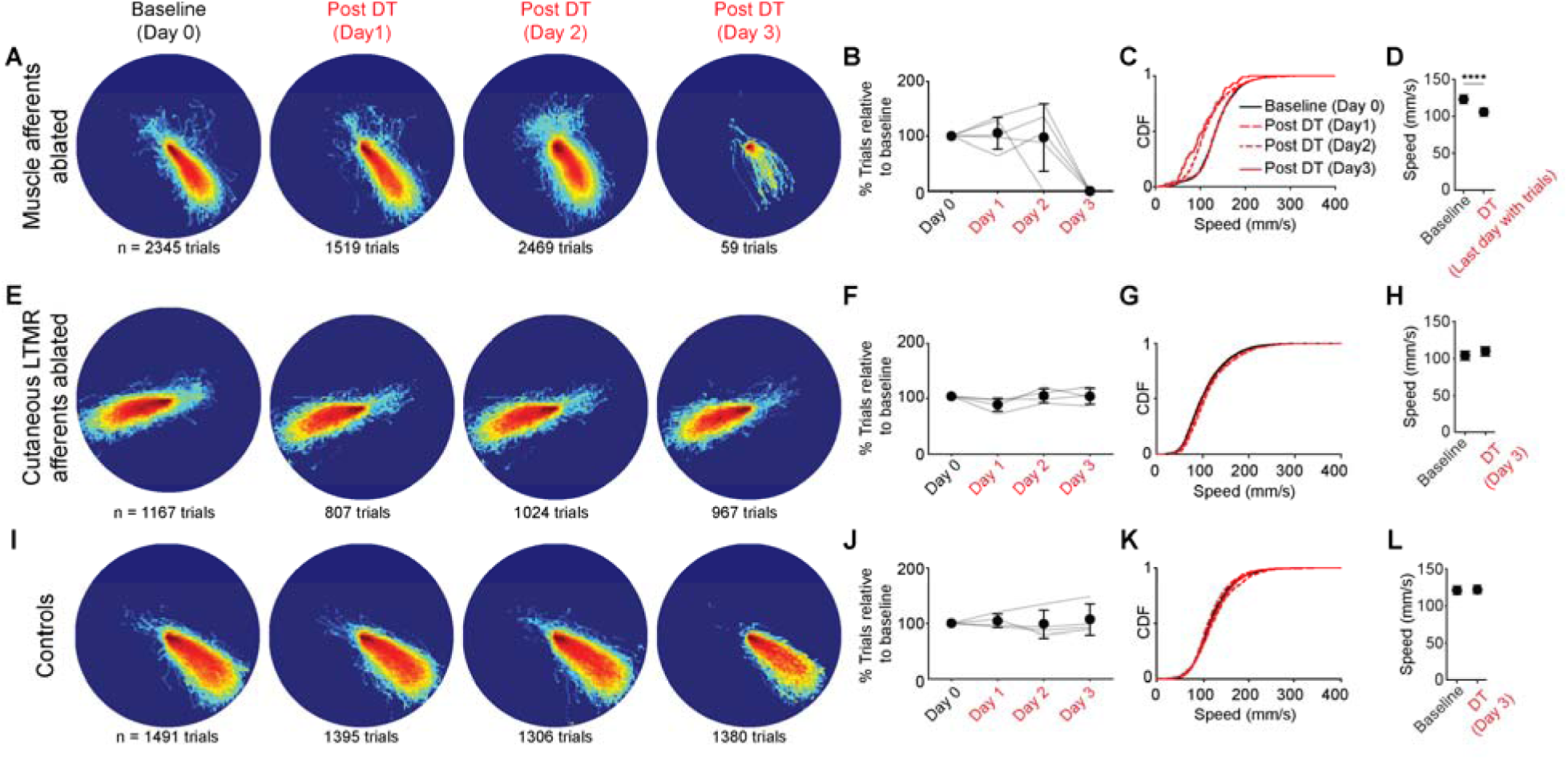
Ablating muscle and tendon afferents blocks the ability to initiate skilled forelimb movements in freely moving mice. **(A)** Heat maps of forelimb reach trajectories in a representative *Pirt::Cre; PV^FlpO^; Tau^ds-DTR^; R26^ds-tdT^*mouse injected with diphtheria toxin (DT) to selective ablate muscle afferents. Forelimb trajectories get progressively disorganized after the day of injection and the animal stops performs the task by day 3. **(B).** *Pirt::Cre; PV^FlpO^; Tau^ds-DTR^; R26^ds-tdT^*injected with DT lose the ability to perform the task by day 3. **(C-D).** Muscle afferent ablation progressively slows down reaching movements across animals. Example distribution of peak reach speed across days in an example animal (C) and means across animals (D). **(E-H).** Plotted as (A-D) for *Avil^CreERT2^; MafA^FlpO^; Tau^ds-DTR^; R26^ds-tdT^* animals injected with DT. This intersection targets a large fraction of the cutaneous LTMR afferents, and their ablation didn’t alter either the spatial structure of the trajectories or the speeds. **(I-L).** Plotted as (A-D) for littermate control animals of those shown in (A-D) that were missing either the Cre or FlpO allele and injected with DT. The spatial structure of the trajectories and the speeds were unaltered by infusion of DT. Values are plotted as median +/- interquartile range. **** denotes a p value<0.0001 for a hierarchical bootstrap test on the medians n = 4 animals.

**Fig. S11.**
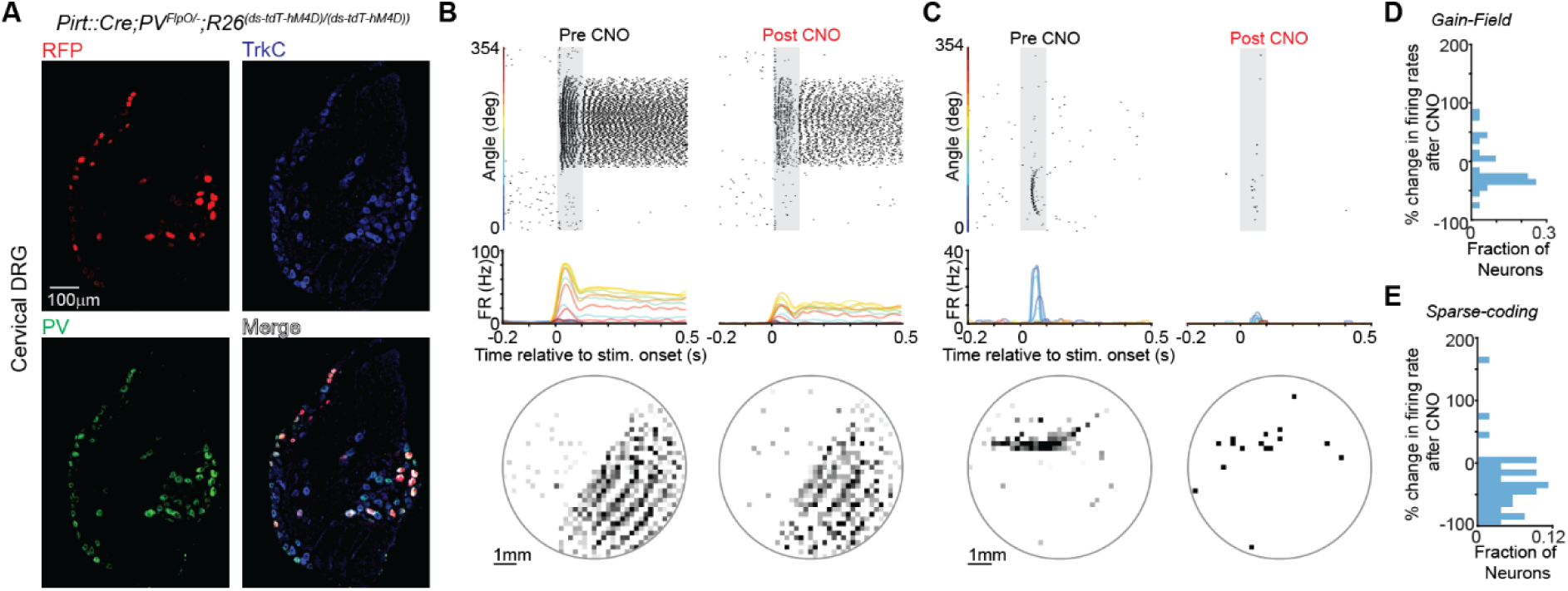
Electrophysiological validation of silencing muscle afferents with hM4D. **(A).** An intersectional strategy to express the DREADD Gi receptor hM4D in muscle afferents. Example images from the dorsal root ganglion of a *Pirt::Cre; PV^FlpO^; R26^ds-tdT-hM4D/ds-tdT-hM4D^*mouse. **(B).** Raster, rate histogram and spatial firing field for an example Gain-field neuron before (left) and after (right) the infusion of CNO in a *Pirt::Cre; PV^FlpO^; R26^ds-tdT-hM4D/ds-tdT-hM4D^* mouse. Note that while the tuning direction is maintained, the peak firing rate is reduced by ∼50% **(C).** Plotted as (B), but for an example Sparse-coding neuron. Note that the infusion of CNO completely disrupted the spatial firing field of this neuron. **(D).** Peak firing rates in Gain-field neurons are reduced by an infusion of CNO **(E).** Plotted as in (D), but for Phasic neurons

**Movie S1. Electrophysiology from the cervical spinal cord of an awake mouse making spontaneous movements.**

Example electrophysiological signals aligned to video and tracked hand kinematics as the mouse makes spontaneous, uninstructed movements. Filmed at 250 frames/second, playback at 30 frames/second.

**Movie S2. Neural geometry for forelimb proprioception.**

Temporal evolution of the neural population trajectories (dPCA) as the limb is moved in different directions.

